# Fifty shades of The Virtual Brain: Converging optimal working points yield biologically plausible electrophysiological and imaging features

**DOI:** 10.1101/2020.03.26.009795

**Authors:** Paul Triebkorn, Jil Meier, Joelle Zimmermann, Leon Stefanovski, Dipanjan Roy, Ana Solodkin, Viktor Jirsa, Gustavo Deco, Michael Breakspear, Michael Schirner, Anthony Randal McIntosh, Petra Ritter

## Abstract

Brain network modeling studies are often limited with respect to the number of data features fitted, although capturing multiple empirical features is important to validate the models’ overall biological plausibility. Here we construct personalized models from multimodal data of 50 healthy individuals (18-80 years) with The Virtual Brain and demonstrate that an individual’s brain has its own converging optimal working point in the parameter space that predicts multiple empirical features in functional magnetic resonance imaging (fMRI) and electroencephalography (EEG). We further show that bimodality in the alpha band power - as an explored novel feature - arises as a function of global coupling and exhibits inter-regional differences depending on the degree. Reliable inter-individual differences with respect to these optimal working points were found that seem to be driven by the individual structural rather than by the functional connectivity. Our results provide the groundwork for future multimodal brain modeling studies.

## Introduction

Brain network simulations allow us to investigate how realistic neural behavior arises from variations in specific biological parameters. Large-scale brain network models can integrate empirical data from multiple modalities into a mathematical framework of complex dynamic systems^1,2^. The Virtual Brain (TVB; thevirtualbrain.org) is a simulation platform that combines local neural mass models (at the level of neural populations) with individual structural connectivity (SC) to simulate whole brain dynamics^3–5^. A unique feature of TVB is the ability to model “personalized virtual brains”, whereby an individual’s diffusion tensor imaging (DTI) derived SC together with functional connectivity data constrains the inter-regional interactions. TVB has previously been used to model biophysical correlates of recovery after stroke^6^, brain activity dynamics pre- and post-surgery in brain tumor patients^7^ and the spreading of seizure activity in epilepsy^8^ to aid the design of therapeutic interventions (see http://www.thevirtualbrain.org for a full list of publications). The global parameters in TVB that determine the spatio-temporal integration across the cortex are conduction speed and long-range global coupling. Conduction speed is the time rate at which the signal travels between the nodes and is used to estimate time delays based on distances derived from an individual’s SC. Global coupling is a scaling factor for the individual’s tractography-derived connectivity weights. Conduction speed and global coupling in TVB and similar models have been associated with important neural mechanisms of healthy and diseased brains^6,9^. Specifically, global coupling has been linked with e.g. tuning the network close to a bifurcation point^10^, exploring the dynamical landscape^9^, motor recovery in stroke^6^, Alzheimer’s disease^11^. Optimal global parameters reproduce many empirical network behaviors such as network synchronization^12^, fluctuations in the system’s stability leading to the emergence of resting-state networks^10^, changes in the attractor landscape^13^, and realistic electroencephalography (EEG) alpha-band activity^14^. Yet, several brain network modeling studies are limited with respect to sample size and the number of metrics fitted - often a single brain activity modality is simulated and a single metric such as functional connectivity (FC; that is the time-averaged correlation matrix based on all regions pairs) is reproduced. However, the assessment across multiple modalities and metrics is essential to validate the models’ overall biological plausibility. In addition, investigating inter-subject variability is critical to disentangle individual from general population aspects.

Both limitations will be addressed in the present study:

1. We utilize a large sample of healthy subjects across the full age spectrum (N = 50, 18 to 80 years of age, mean 41.12±18.20; 30 females and 20 males) for “virtualizing” their brains and simulating individual whole-brain neural and functional magnetic resonance imaging (fMRI) blood oxygen dependent (BOLD) signals.
2. We aim to show that there exists a specific set of neural parameters for which virtual brains operate in a biologically realistic manner in the resting-state, and that these parameters converge to a unique point for each individual across several metrics. We focus on the global parameters of conduction speed and long-range coupling.

Specifically, we demonstrate that the parameter subset for this optimal fit converges for the following metrics:

a. correlation between empirical and simulated fMRI BOLD FC;
b. fit of simulated to the empirical fMRI BOLD FC dynamics (FCD), that is the switching of functional connectivity over time^15^;
c. frequency of the mean neural signal with a focus on the most prominent human EEG feature that is the alpha (10Hz) rhythm^16^ during wakefulness and delta (2Hz) during sleep^17^ and its known multistable power distribution^18^;

We aim to demonstrate not only feasibility but also specificity of individual model predictions and to explore the mechanisms that lead to the emergence of the empirical data features under investigation – with a special focus on the bistable electrophysiological alpha rhythm in a full brain network context advancing our previous work lacking this aspect^18–20^.

## Results

The best predictions of empirical signal features (FC, oscillations and bimodality of the neural signal) arise as a function of particular ranges of neural parameters (**Fig. 1**), which we term the optimized parameters. Empirical features were reproduced at a wide range of conduction speeds, but only in a narrow range of global coupling values (**Fig. 1**). For the delta (alpha) set, these optimal parameters were around *G*∈*[0.1,0.14] (G*∈*[0.0280,0.031]),* where *G* is the global coupling. The mean best correlation to empirical data for individual subjects was 0.45 (0.514) and the highest correlation 0.66 (0.68) in the delta (alpha) set. Bimodality was found at *0.11<G<0.14* (*0.0286<G<0.031*) and was more likely to appear at higher conduction speeds in the delta (alpha) set (**Fig. 1B**). The fastest average oscillation was observed at lower couplings than the peaks of the two other metrics (**Fig. 1C**). With increasing global coupling, an increase in frequency was observed in the delta set, which continued until a critical point where the oscillation speed began to decrease again (**Fig. 1C, left**). In the alpha set, the frequency is continuously decreasing as a function of global coupling (**Fig. 1C, right**).

**Figure 1.**
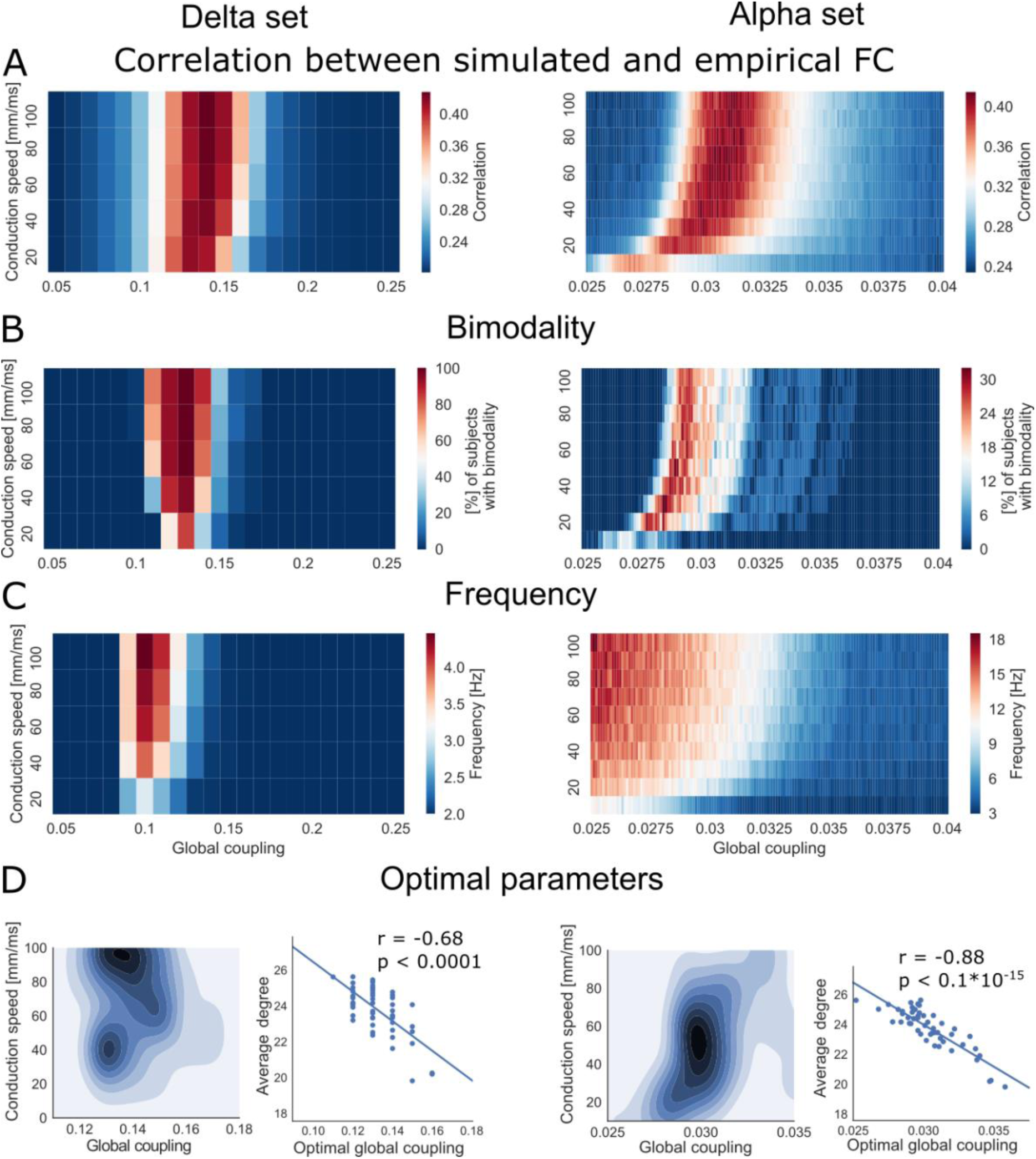
Overview of the parameter exploration generating oscillations in the alpha and delta rhythm range. Both simulation sets (delta set on the left and alpha set on the right) varied the parameters conduction speed and global coupling, but across different ranges. **A**: Prediction quality (comparison of simulated and empirical data via means of FC correlation) of all simulations averaged across all subjects. **B**: Bimodality assessed on the mean neural signal using Hartigan’s dip test. There is an overlap between best empirical-simulated FC fit and bimodality of the neural signal at distinct global coupling (compare red-colored areas in 1st and 2nd rows) for both simulation sets. **C**: Frequency of the oscillating mean neural signal. **D**: Distribution of optimal parameter points for all subjects derived by correlation to empirical FC. Density plot was smoothed with a Gaussian kernel. Best fits can be seen across a wide range of speed, but only across a few coupling values. Some subjects had best fits at conduction speeds around 40mm/ms, whereas others best fit around 100mm/ms in the delta set. The scatter plots show a linear relation between optimal global coupling (assessed by maximum correlation empirical-to-simulated FC fit) and average degree (defined as the mean of row sums in each subject’s SC matrix) (delta: r=-0.68, p<0.0001, alpha: r=- 0.88, p<0.1·10^-15^).

Because averaging over subjects obscures individual features, we also extracted an optimal parameter set for each subject defined as the best fit to empirical FC (density plot in **Fig. 1D**). Again, we observed that best fits to empirical data were distributed across many conduction speeds but in a narrow range of global coupling. Interestingly, there were two modes of highest fit in the delta set along the conduction speed axis, suggesting a large degree of between-subject variability. Within the alpha set, the optimal parameters were distributed around one single mode. Distribution parameters of individual optimal points are listed in **Supplementary Table 3**. We found no significant differences in optimal working point distributions between genders (**Supplementary Table 5**). We also confirmed that optimal global coupling correlates with the average degree of the SC (i.e. average number of connections of a brain region, **Fig. 1D**)^28^.

### Empirical-to-Simulated FC Fit: Subject Specificity

We were interested in the subject specificity of a subject’s simulations, and the degree to which parameters are driven by SC and empirical FC. Therefore, we assessed how a subject’s set of simulated FCs fits to empirical FC of other subjects. This procedure yielded one highest possible correlation for each pair of subject’s sets of simulations and empirical FCs (‘highest correlation matrix’, **Fig. 2A**). There was no individual subject fit of simulated to empirical FCs (**Fig. 2B, left**). However, there was an individual subject fit of the SC to the simulated FC (stronger correlations on the diagonal of **Fig. 2B, middle**). This finding is also shown on the density distributions, where individual fits (SC and simulated FC come from the same subjects) outweigh all-to-all fits (where SC and simulated FC come from different subjects, **Fig. 2B, right**). The results are shown for the alpha set but resemble those for the delta set (**Supplementary** Fig. 10). We also examined variations between subjects’ SCs and FCs (**Supplementary** Fig. 10).

**Figure 2.**
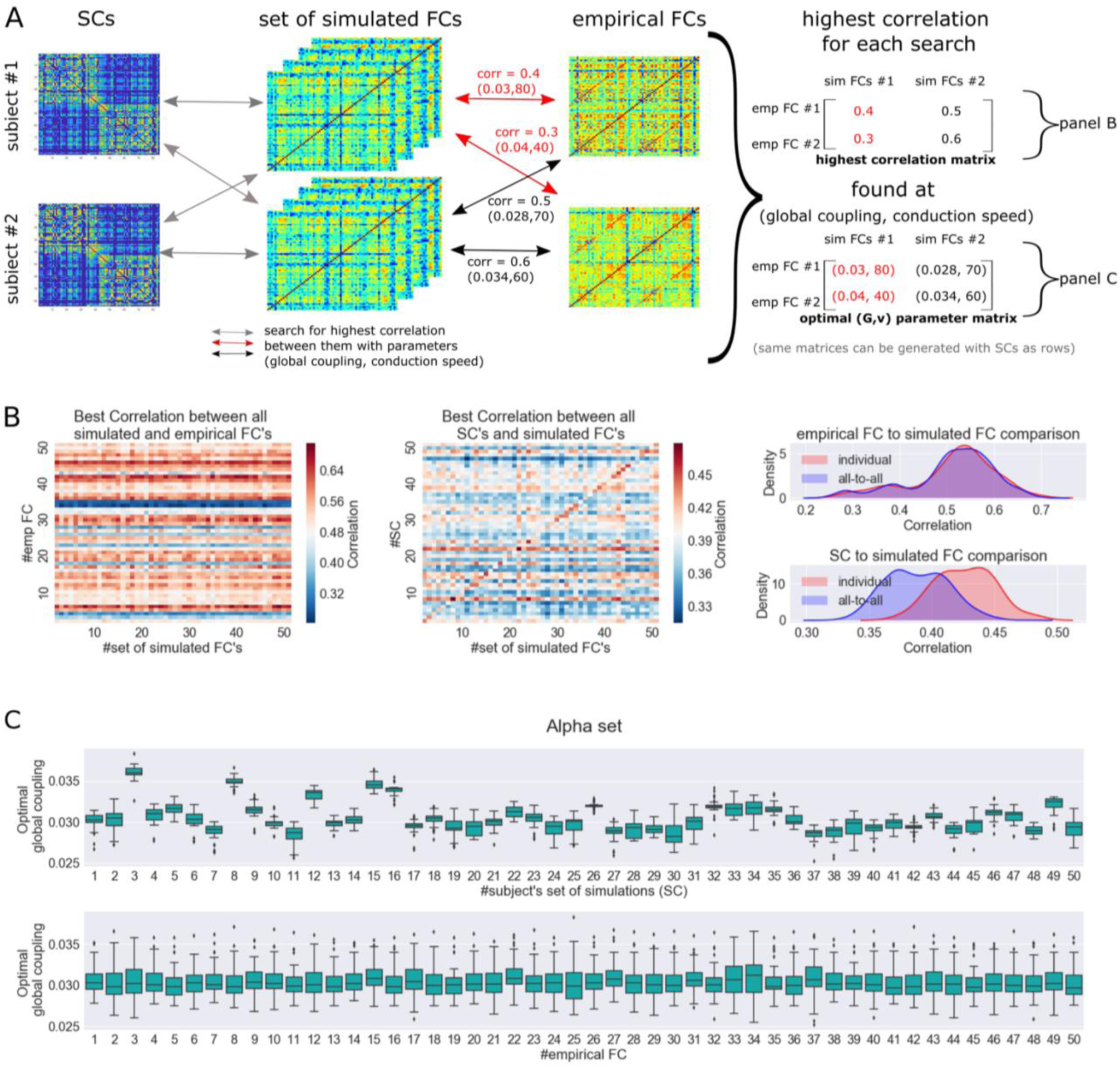
Cross-subject comparison of simulations results. **A**: Comparing all simulations of one subject to their own empirical FC yields one optimal parameter combination. However, this parameter combination does not have to be optimal in terms of goodness of fit if we compare the same set of simulations to other subjects’ empirical FC of other subjects. Performing this analysis for all 50 subjects yields two 50×50 matrices: the optimal parameter combinations, further denoted as ‘optimal (G,v) parameter matrix’ and the ‘highest correlation matrix’. The schematic shows the computation of those two matrices for two example subjects (subject #1 and subject #2). The rows of these matrices represent the subject’s empirical FC compared to the set of simulated FCs based on the subject number equal to the column number (comparisons visualized as red and black arrows). Both matrices can also be calculated based on comparisons of the SC of each subject (as rows) with the set of simulated FCs based on the corresponding subject number of the column (grey arrows in the schematic). **B**: Alpha set: Left: We correlated all 1510 simulations of one subject to all 50 empirical FCs. For each empirical FC that we correlated to, there was one best (i.e. highest) correlation. The entry (i,j), 1≤ *i*, *j* ≤50, in then displayed ‘highest correlation matrix’ represents the highest correlation based on the comparison of the empirical FC matrix of subject i with the set of all simulated FCs of subject j. Along the diagonal of the matrix, we found the highest possible correlation for the individual subject prediction (i.e. predicting a subject’s empirical FC by using the same subject’s SC for simulation). Horizontal red and blue stripes already indicate that some empirical FCs can be predicted better than others, regardless of the SC used for the simulation. Middle: The same as the left matrix but now comparing set of simulated FCs to SCs, instead of empirical FCs. We tested the possible prediction of the SCs from the simulation. A vague red diagonal indicates a best individual fit, pointing to the fact that the simulated FC contains typical features of the used SC. Right: To test for subject specificity, we took the diagonal values (i.e. subject’s individual prediction) and tested their distribution against the values above and below the diagonal (i.e. all-to-all prediction). Comparing simulations to empirical FCs showed no significantly better individual prediction. Individual SCs can be predicted significantly better than all-to-all prediction (two-sample Kolmogorov-Smirnov test d = 0.4963 (d = 0.5492) and p < 0.1·10^-9^ (p < 0.1·10^-12^) for delta (alpha)). This result indicates how strongly the SC shapes the simulated FC. The density plot was smoothed with a Gaussian kernel. **C**: Optimal parameter combinations are subject-specific (results displayed for the alpha set): We analyze how the optimal parameter distributions change between subjects. The parameters are influenced by the underlying SC as visible in the upper panel. Each boxplot displays the optimal global coupling values for each subject (best reproducing empirical FCs of all subjects). The variance of optimal global coupling for a given subject’s SC is small compared to the variance of the same across individual subjects’ SCs. In the lower panel, we display the influence of the empirical FC on the distributions of optimal global coupling values. In contrast to SC that strongly determines the value of optimal global coupling, the individual empirical FC does not determine the value of optimal global coupling to the same degree. This result underscores the high importance of the SC for determining optimal coupling parameters.

The simulated FC that fits best to the same subject’s empirical FC may not necessarily be the same simulated FC that fits best to other subjects (i.e. optimal parameter points may vary if we compare a set of simulations of one subject to empirical FCs of other subjects). Therefore, we extracted the corresponding optimal parameter combinations (‘optimal (G,v) parameter matrix’, **Fig. 2A**). The results show that optimal coupling is mostly determined by the underlying SC (**Fig. 2C**), which is strengthened by eta-squared effect sizes (**Supplementary Table 4**). The effect is more visible in the alpha set with an eta squared of 0.7968 for coupling explained by SC compared to 0.0175 by empirical FC. The delta set shows the same characteristics with small differences in effect sizes. However, conduction speed is neither determined by the SC nor by empirical FC (**Supplementary Table 4 and Supplementary** Figure 15). The change in simulation result is very small for different conduction speeds. Thus, a variety of transmission speeds can be seen as optimal.

### Mechanisms Underlying Electrophysiological Alpha Rhythm Bimodality

To understand the mechanisms underlying electrophysiological rhythm bimodality, we conducted further simulations where we decreased global coupling gradually in five steps to zero (**Fig. 3** and **Supplementary Video 1**). At *G = 0*, the mean field potential was the time series generated solely by the local neuronal population model. We observed a bursting behavior in the neural signal that has been documented by the authors of the population model^34^. A fold/homoclinic bifurcation is characteristic for this type of burst, where the trajectory follows a limit cycle. Meanwhile, the slow variable (*z*) increases and moves an unstable fixed point towards the limit cycle, generating a saddle homoclinic bifurcation, i.e., the unstable fixed point touches the cycle and breaks it up. The trajectory then switches towards a stable fixed point, which corresponds to the resting period. During “rest”, the slow variable decreases and moves the unstable system back towards the stable fixed point creating a fold bifurcation - a transition from rest back to spiking. This bifurcation type has been described for the single Hindmarsh-Rose neuron^30^, the foundation of the here used population model.

**Figure 3.**
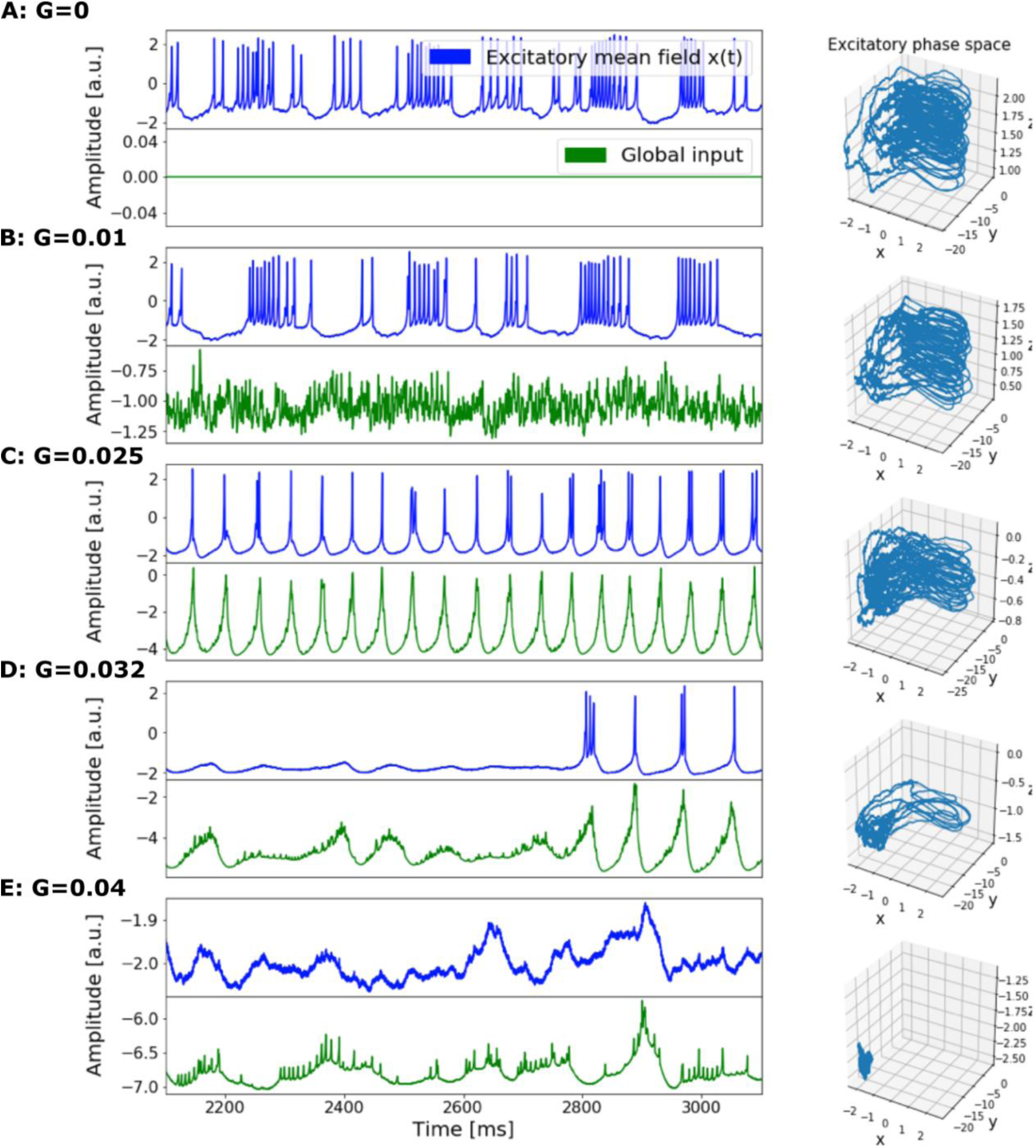
Bimodal behavior of electrophysiological neural dynamics in the brain network model. Time series represent simulated neural activity in the left inferior temporal cortex of an example subject. Local parameters were chosen as in the alpha set (Table 1, Methods) with conduction speed set to 100 mm/ms. Plots from top to bottom with increasing global coupling A: G=0, B: G=0.01, C: G=0.025, D: G=0.032 and E: G=0.04. Blue time series depict the mean field potential (i.e. neural signal). The global input to the inferior temporal cortex from all other nodes, scaled by connectivity weights and global coupling, is colored in green. On the right, the excitatory phase space is depicted with excitatory state variables x, y and z. A: For G = 0, it shows the switching between a limit cycle and a fixed point, creating a burst-like behavior. B: As we increased global coupling (to G = 0.01), regions were unsynchronized and the global input consisted of noisy fluctuations. C: For even higher couplings (G = 0.025), we observed a synchronization of local dynamics with global input. The regional neural signal displayed single bursts with few spikes. There were only a few turns on the limit cycle in the phase space before the homoclinic bifurcation occurs. The global input became even more negative (i.e. -4). We show that the spiking in the regional neural signal coincided with the times at which the global input reached zero. This result can be interpreted as a temporary reduction in global inhibition that leads to spikes in the local model. D: The state of bimodality occurred in-between the synchronized and the oscillation death state (G = 0.032). Namely, we observed a switching between oscillation death and spikes that occurs in synchrony with a reduction in global inhibition. E: For very high global coupling (G = 0.04), oscillation death occurred and very slow noisy fluctuations in the neural signal were observed. The neural signal and the global input had negative amplitudes and fluctuated very little. The trajectory remained at a stable fixed point. We show this behavior also in **Supplementary Video 1.**

**Table 1:** Global and local parameters for delta and alpha set simulations.

|  | parameters | delta set | alpha set |
| --- | --- | --- | --- |
| **Global parameters** | *global coupling:* | linear | linear |
|  | G (multiplicative) | [0.05, 0.25], step size: 0.01 | [0.025, 0.04], step size: 0.0001 |
|  | conduction speed [mm/ms] | [20, 100], step size: 20 | [10, 100], step size: 10 |
|  | *integrator:* | HeunStochastic | HeunStochastic |
|  | integration step size | dt = 0.01220703125ms | dt = 0.05ms |
|  | additive noise | D = 1.0 | D = 0.001 |
|  | *monitors:* |  |  |
|  | <u>SubSample</u> (of neural signal)<br>sampling period | 3.90625ms (= 256Hz) | 3.90625ms (= 256Hz) |
|  | <u>BOLD</u><br>HRF Kernel<br>sampling period | MixtureOfGammas<br>500ms (= 2Hz) | MixtureOfGammas<br>500ms (= 2Hz) |

|  | duration of simulated time series | 180s | 300s |
| --- | --- | --- | --- |
| **Local parameters** | r | 0.006 | 0.006 |
|  | a | 1 | 1 |
|  | b | 3 | 3 |
|  | c | 1 | 1 |
|  | d | 5 | 5 |
|  | s | 4 | 4 |
|  | x<sub>0</sub> | -1.6 | -1.6 |
|  | K<sub>11</sub> | 0.5 | 4 |
|  | K<sub>12</sub> | 0.1 | 1.6 |
|  | K<sub>21</sub> | 0.15 | 0.15 |
| | $\sigma$ | 0.3 | 0.4 |
| | $\mu$ | 2.2 | 2.2 |

With increasing G, the system becomes increasingly shifted towards the fixed point, i.e., a point that once the system enters no changes in the dynamics occur anymore. It reaches oscillation death at G = 0.04. The global input range was negative (y-axis of **Fig. 3B)** because the neuronal population model’s excitatory state variable x (displayed) has a negative baseline (and positive values during spikes). That is, in the here used population model, an excitatory population *decreases or increases* the membrane potential of another neural population, depending on the current membrane potential. For G=0.032, we observe a bimodal state between oscillation death (i.e. residing in the fixed point) and oscillations (i.e. limit cycling, **Fig. 3D**).

We also analyzed bimodality and the frequency of the neural signal of each region individually (**Supplementary** Fig. 11-12). Similar to what we found for the global signal, regional signals at critical values of conduction speed and global coupling also shape oscillations and bimodality. Furthermore, regional distribution of the bimodal power distribution was strongly determined by the SC for the delta set, namely a strong degree, that is node connectivity, was linked to the emergence of bimodality (**Supplementary** Fig. 12). For the alpha set, nearly all regions demonstrated bimodal power distribution at the optimal parameter point. This difference is caused by local parameters (see e.g. **Supplementary** Fig. 7).

### Functional Connectivity Dynamics

**Fig. 4** shows empirical and a simulated FCD for one example subject. Though the highest correlations of subsequent FC matrices in our simulations are lower than in the empirical data, persistent (over time) FC patterns in the BOLD signal can be observed for both empirical and simulated time series (**Fig. 4A**). Interestingly, the optimal range of global coupling for the FCD fit, bimodality, and static FC fit are highly overlapping (**Fig. 4B**). Thus, only when our local model presents bimodal behavior, switching of modes is generated in our global network. **Supplementary Video 2** shows simulated and empirical FCDs and FCs for this example subject while sliding across the range of global coupling values.

**Figure 4.**
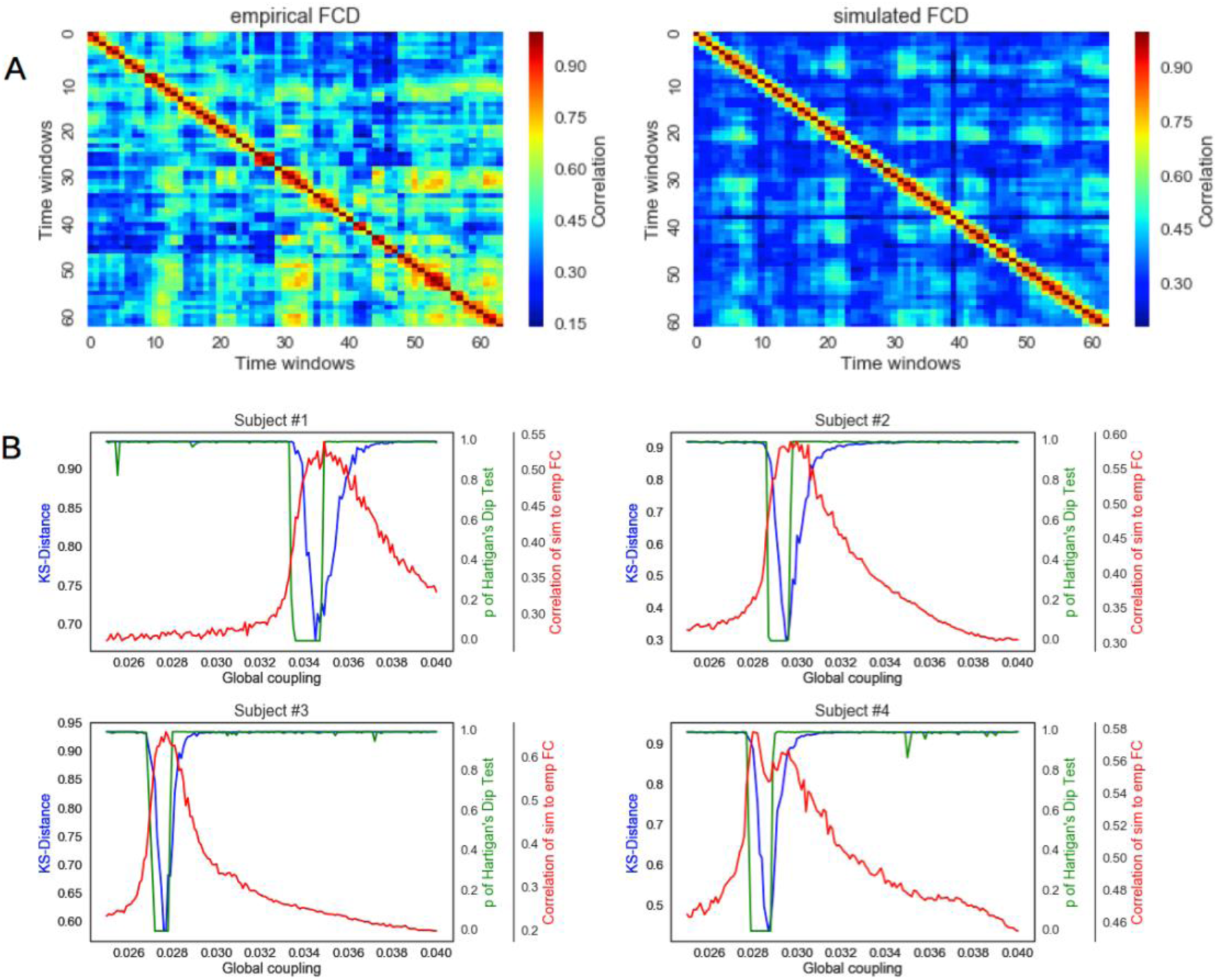
Functional connectivity dynamics align with bimodal behavior of neural dynamics. **A:** Example of empirical (left) and simulated (right) FCD of an example subject (female, 47 years old). **B:** Metrics of 22 minutes simulations for four sample subjects (sex/age: f/30, f/47, m/20, m/73). In blue, the Kolmogorov-Smirnov distance between the simulated and empirical FCD. Best FCD fit (i.e. lowest Kolmogorov-Smirnov distance) occurs at the same range of global coupling as bimodality of the global signal (green) and best fit between FCs (red).

## Discussion

In this study, we conducted a large parameter space exploration with TVB, simulating BOLD fMRI and electrophysiological neural signals for 50 individual human brains. Minute variations in global coupling and conduction speed were shown to give rise to fluctuations in 1) the correlation between empirical and simulated BOLD FC, 2) frequency of the mean neural signal, 3) the bimodality in the frequency power in the alpha band, as well as fluctuations in 4) the fit of FCD. We have shown that all of these features converge at a particular parameter point unique to each subject and demonstrated that there are reliable inter-individual differences in biophysical parameters that lead to optimal working points.

### Optimal Parameter Values

Our range of explored conduction velocities (10-100m/s) was comparable to that found in the adult primate brain (5-94m/s), where neural axons are heavily biased towards those with large diameters and high conduction speeds^21^. Similarly, we found that our empirical signal features were best reproduced at conduction velocities higher than 20mm/ms with negligible differences above this threshold. Optimal global coupling is largely dependent on the underlying SC characteristics^22^. Our global coupling factor was quite low, and comparable to studies with dense SCs due to coarse parcellation schemes (optimal *G*: delta set H0.13, alpha set H0.03). Previous results for global coupling were ∼0.007^23^, ∼0.016^24^ and ∼0.1^25^. The found differences in optimal global coupling parameters for the delta vs. alpha set are in line with a previous study applying a Wilson-Cowan model^26,27^.

We found that the ability of the model to approximate the empirical neural signal was determined predominantly by the global coupling, and less by the conduction speed. Previous resting-state models have emphasized the critical role of the inter-nodal coupling in determining spatial structure^23,28,29^. Spiking models, such as the reduced Wong-Wang^30^, in contrast to simple oscillators, may not necessitate time delays, as synchronization is decreased through system heterogeneity, noise, and greater time between individual spikes^31^. Moreover, delays have a greater influence on dynamics in directed (asymmetric) than undirected (symmetric) networks^22^, such as our DTI-inferred SCs. This result arises because symmetric matrices with real entries have real eigenvalues. Thus, the connectivity affects the system’s equilibrium but not its dynamic oscillatory behavior^22^. The Hindmarsh-Rose model forming the basis of the current study is similar to the FitzHugh-Nagumo, where time delays were not critical in shaping network dynamics^32–34^.

The observed realistic properties of the simulated neural signal do not arise linearly with global coupling, but rather slowly emerge, sharply peak, and drop off afterwards. A number of other resting-state neural network properties, such as system multistability^10^, cluster synchronization^23^, emergence of resting-state networks^28^, instability of the equilibrium state^22^, or system efficiency^35^ have similarly been shown to develop as a function of this inter-regional coupling and/or conduction speed. Similar to what we noticed in our study, these behaviors emerge at the point where global coupling produces the highest correlation between empirical and simulated FC. Taken together, these findings point to the significance of a critical point at which the system operates optimally.

### Convergence of optimal parameters for different experimental features

We showed that the region of parameter space that gives rise to the best fit between simulated and empirical (static) FC, and bimodality, coincides with the best model of FCD. Our simulations reproduced time-varying features of the signal relevant to capturing individual differences^36^, and other resting-state phenomena^15^. Our results are in line with previous findings showing that the meta-stable state is a function of a bifurcation parameter along with global coupling, and, importantly, that the best FCD fit is achieved at this meta-stable state^15^.

### Reproducing EEG features

We found that global coupling which exceeded a certain threshold led to a sharp reduction in oscillation frequency resulting in a predominantly delta frequency range with low amplitude. Further, we were able to reproduce bistable switching between two distinct alpha modes, a feature of empirical EEG^19^. Transmission delays and global coupling were previously shown to affect the mean power when simulating magnetoencephalography signals with brain networks, and the best model fit was found in the alpha band power^31^. However, this work is limited to the traditional mean power metric, without taking into account the bistable switching between two distinct alpha modes that occurs in empirical neural signal. The largely overlooked multistable nature of these neural processes should be taken into account^26^. Converging evidence indicates that the different alpha modes are linked to cognitive processing^37^. We show that this bimodality, which has previously been modelled as a function of noise^18^, arises also as a function of optimal global coupling. This result can be contrasted with a previous study^18^, where bimodality was modelled on the neural signal of a single node, thus global parameters would likely not play a large role in determining its behavior.

As a sub-analysis of the newly added fitting criterion of bimodality, we sought to elucidate the mechanisms behind the emergence of bimodality in our model. For low values of global coupling (i.e. *G=0*), we observed bursting behavior at the local model. Such square-wave bursting behavior, as well as single spikes and chaotic behavior have been observed previously in the here used population model and the underlying single-neuron Hindmarsh-Rose model^32,34^. In the current population model, the activity of excitatory neurons is transmitted into the global network, which corresponds to a biological global excitatory coupling in the brain. The baseline of this activity is around -2[a.u.] corresponding to a neuron at rest and transmitting global inhibition, while spikes peak at +2[a.u.]. Thus, at low values of global coupling, bursts/spikes still occur at a node as global inhibition is not strong enough to suppress this activity. Increasing global coupling yet further, results in oscillation death because global inhibition is strong enough to cancel out all activity. In between these two scenarios, there is a narrow range of coupling values where a switching between periods of spiking behavior and periods of oscillation death can be observed, giving rise to a bimodal power distribution. Taking into account that different nodes have different degrees, we can explain that some nodes reach bimodality and oscillation death earlier than others (receiving more global input than others). Thus, inter-regional input plays an important role in driving oscillatory activity, as seen previously in the alpha band^31^.

### Inter-individual differences

We show that there are inter-individual differences in the optimal long-range global coupling. This result suggests that the optimal point of functioning is highly dependent on the individual’s anatomical architecture in combination with the spatio-temporal dynamics of the model. Our findings indicate the importance of conducting parameter space explorations at the individual level, as more standard group-level analyses may obscure important and reliable variability. Moreover, we show that individual variability in the connectomes was driven primarily by the simulated FC and the underlying SC, and to a lesser extent by the empirical FC, which is important for exploring subject-specific parameters. Thus, the best starting point for optimal global coupling can be drawn from the SC rather than the empirical FC. The same conclusion cannot be drawn for conduction speed. There may be other parameters that we did not explore which may be more determined by the empirical FC than the SC.

The observed simulated-to-empirical FC correlations (delta set: mean *r*=0.45; max *r*=0.66; alpha set: mean *r*=0.51, max *r*=0.68) are in the range of those obtained in parameter explorations with similar models (*r*=0.427^28^; *r*=0.42^13^; *r*Η0.4^10^). The ability of our model to reproduce the empirical data is limited by the quality of current diffusion techniques. Existing links that are not captured by the SC may conceal how real structural connections influence FC. However, in light of these inherent limitations, our model has captured a number of important properties of the neural and BOLD signal. Another limitation of the current study is the computationally expensive full-grid search for the optimal parameters, which should be improved with e.g. Bayesian optimization techniques in the future^38^.

In sum, we have demonstrated that as global coupling and conduction speed approach an optimal working point, the behavior of the neural system reproduces the resting-state BOLD static and dynamic FC, as well as the neural signal. With the present work, we aim to provide guidance for personalized brain network modelling with the neuroinformatics platform TVB. A systematic exploration of other population models and comparisons of their respective parameter landscapes will help to further test and validate the biological interpretability of parameters and model variables.

## Methods

### Data Acquisition

Simultaneous EEG-fMRI data, diffusion-weighted MRI (dwMRI), and structural MRI were recorded from 50 healthy adult subjects at Berlin Center for Advanced Imaging, Charité University Medicine, Berlin, Germany (18 to 80 years of age, mean 41.12±18.20; 30 females and 20 males, for the distributions see **Supplementary** Fig. 14). Subjects were voluntarily recruited, and had no self-reported history of cognitive, neurological or psychiatric conditions. Informed consent was provided to subjects before participating in the study, and the research was conducted in accordance with the Code of Ethics of the World Medical Association Declaration of Helsinki and approved by the local ethics community at Charité University. The data acquisition procedures are those reported by our colleagues^5^ and are summarized here.

Images were acquired on a Siemens Tim Trio MR scanner (12-channel Siemens head coil). Anatomical and dwMRI acquisition was performed first, after which subjects were taken out of the scanner to have their EEG cap set up. Subjects then returned to the scanner for simultaneous EEG-fMRI measurements. Subjects laid in the scanner with their eyes closed and were asked not to fall asleep.

High resolution 1×1×1mm T1-weighted scans were acquired with an MPRAGE sequence: repetition time (TR) =1900 ms, echo time (TE) =2.25 ms, flip angle (FA) =98, field of view (FoV) =256 mm, 256 matrix, 192 sagittal slices, slice thickness =1mm. EEG and fMRI were recorded for 22 minutes at resting state. We used an echo planar imaging (EPI) T2* sequence: 666 volumes, TR= 1940ms, TE=30ms, FA=788, FoV=192, 64 matrix, voxel size =3×3×3mm^3^, 32 transversal slices, slice thickness =3mm. The first five images of the BOLD fMRI series were removed to prevent disturbance from saturation effects. Diffusion-weighted MR echo-planar measurements were conducted with TR=7500ms, TE=86ms, FoV=220mm, 96 matrix, voxel size =2.3×2.3×2.3mm, 61 transversal slices, slice thickness =2mm; 64 diffusion gradient directions with b-values =1000s/mm^2^.

### Brain Imaging Data Preprocessing

Preprocessing of anatomical, diffusion and functional data was performed using our pipeline for derivation of individual connectomes^39^. The preprocessing steps are described in detail there, and more briefly below. Preprocessing of T1-weighted images included the following: motion correction, intensity normalization, extraction of non-brain tissue, brain mask generation, and segmentation of cortical and subcortical grey matter. We used the parcellation from the Desikan-Killiany Atlas^40^ implemented in FREESURFER to divide the cortex into 68 regions of interest. A quality check of the parcellation of individual high-resolution T1- weighted scans was performed manually.

The dwMRI data was preprocessed as follows: motion and eddy current correction, linear registration of b0 image to individual T1-weighted image. The anatomical cortical parcellations were transformed to individual diffusion space, where probabilistic tractography was conducted. Spherical deconvolution was applied to constrain tractography, using MRTrix streaming method, which has the ability to identify crossing fibers (FA threshold =0.1)^41^. The gray matter – white matter interface was exhaustively sampled, streamlining up to 200,000 times from each voxel (radius of curvature =1mm, maximum length =300mm). Binary connections for each region pair, based on the Desikan-Killiany atlas, were then aggregated, and a 68-by-68 SC was obtained. The region labels are listed in **Supplementary Table 1**. A weights and distance matrix were calculated for each subject. Weights matrices were the number of voxel pairs between two ROIs with at least one track found. We modified the weights matrix by using the common logarithm of its values and then normalizing to its highest value. Distance matrices were the mean track length of fibers between each pair of ROIs, measured in mm.

In brief, the functional data was preprocessed as follows: motion correction, brain extraction, high-pass filter (100s), registration to individual T1-weighted anatomical scans to apply high-resolution parcellation mask to the fMRI. Additionally, quality of BOLD signal was analyzed with temporal signal to noise to maps^42^. BOLD signal was averaged across voxels within brain regions. Functional connectivity was computed using a Pearson’s linear correlation coefficient on the BOLD data of each region pair.

### Global Dynamics of TVB Model

The global behavior of the system is driven by interactions between individual nodes, as described by the conduction speed and long-range coupling. Conduction speed is used to estimate time delays based on distances from DTI data. Thus, propagation of neural signal is faster for dense, myelinated neural fibers compared to smaller fibers for the same distance^43^. With a fixed distance, a large conduction speed results in a short time delay; a small conduction speed leads to a large time delay and slower system. The global dynamics at a node *i* can be described by

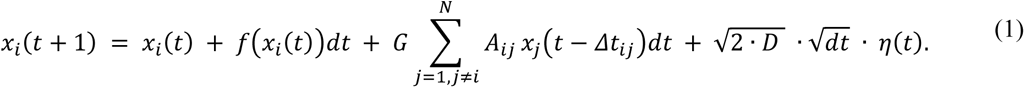

The mean-field potential of node *i* is integrated across time *t={1, …, T}* by taking the sum of its current potential *x_i_(t)*, the derivative of the local model *f(x_i_(t))dt*, the scaled input of all other nodes *j={1, …, N}, jī*, and the influence of noise. The parameter *D* scales the dispersion of the Gaussian distribution *η(t)* from which random values are drawn. Connectivity weights (i.e. individual diffusion-tractography-derived SC weights) are defined here as *A_ij_*, where the connection weight scales the influence of node *i* on node *j*. The time delays of signal propagation, *Δt_ij_=d_ij_/v,* are influenced by distances *d_ij_* (i.e. diffusion-tractography-derived distances between regions) and conduction speed *v*. The long-range coupling factor *G* provides an additional scaling of influence.

The simulations were performed with TVB v1.5. TVB input was the individual SC, and a combination of local and global parameters describing the nodal and network-level dynamics. Different parameter combinations were simulated in two sets for each individual subject, with global and local parameters described in **Table 1**. The neural signal in the first set of simulations oscillated with a frequency of around 2- 5Hz and around 8-12Hz in the second set. We call them delta and alpha set, respectively. Because we were interested in the bimodality mechanisms in the alpha, we conducted more extensive explorations of the alpha set.

The different simulations produced one 3D matrix with the dimensions [subjects, conduction speed values, global coupling values] for each simulation set. The elements of the matrix are correlation coefficients which quantify the similarity between the empirical and the simulation BOLD signal (empirical-to-simulated FC fit). Similarly, a 3D matrix was calculated for bimodality and frequency analysis of the mean neural signal. To visualize our results, we averaged over the [subjects] dimension of each 3D matrix. The result is a 2D matrix with dimensions [conduction speed and global coupling] whose elements are the averaged correlation coefficients over all subjects, the percentage of subjects presenting bimodality in their power distribution and the averaged frequency (in Hz) over all subjects.

All computations described in the following section were performed on a high-performance computer consisting of multicore CPUs at a clock rate of Η2.50 GHz per CPU core and 8 GB RAM per core. The time per simulation was approximately 10h (1.6h) for integration step size 0.01220703125ms (0.05ms) for the delta (alpha) set. In total, we tested 1615 parameter combinations for each subject resulting in 80750 simulations with a total simulation length of Η6554h. The total computation time was Η230000h. Simulation scripts including optimal parameter values and imaging-derived data used to constrain the models are made available via EBRAINS – Human Brain Project’s European Brain Research Infrastructures.

We coupled nodes via long-range connections in the model. The anatomical connectome served as the structural backbone of these connections. In both sets, we varied two free parameters to examine the spatiotemporal dynamics: the global coupling *G*, which scales the input to a node from others, and the conduction speed *v* (mm/ms), which represents delays in the propagation of the signal. The global coupling scaling factor *G* is named “a” in the TVB interface. In this article, we refer to it as global coupling or “G”. In the first simulation set (delta set), conduction speed was explored in a range from 20mm/ms to 100mm/ms in steps of 20mm/ms and global coupling from 0.05 to 0.25 in steps of 0.01 (**Table 1**). This procedure gave rise to 5250 simulations (50 subjects x 105 parameter combinations). For the second set (alpha set), achieving bimodality in the alpha frequency oscillations, we varied conduction speed from 10mm/ms to 100mm/ms in steps of 10mm/ms and global coupling from 0.025 to 0.04 in steps of 0.0001, for a total of 75500 simulations (50 subjects x 1510 parameter combination, **Table 1**). Both sampling rates are much faster than the Nyquist rate for the frequency band of interest, allowing both delta and alpha rhythms to be resolved in the signal. The integration step sizes were then chosen to allow fast computation and guarantee stability.

### Local Stefanescu-Jirsa 3D Model

We used a local Stefanescu-Jirsa Hindmarsh-Rose 3D (SJHMR3D) mean-field model^34^ in TVB to simulate nodal neural behavior. The SJHMR3D is a biologically realistic model based on a network of heterogeneous excitatory and inhibitory coupled Hindmarsh-Rose neurons^33^ that exhibit a complex repertoire of neural behavior - such as synchronization and random firing, multi-clustering, bursting, transient oscillations and oscillation death. Coupled neurons of the detailed network showed reorganizing behavior into clusters with similar dynamics. Using mode decomposition techniques, the authors in^33^ were able to capture cluster dynamics from the detailed network in a low-dimensional representation of only three modes. The set of equations in **Eq. 2** describes local dynamics at a node *i,* with each state variable being a vector of length three to represent the different modes. The SJHMR3D model aims to describe the average membrane potential of a population of neurons that are electrically coupled. The model is comprised of six differential equations representing interconnected neural populations of excitatory *(x, y, z)*, and inhibitory *(w, v, u)* populations as such

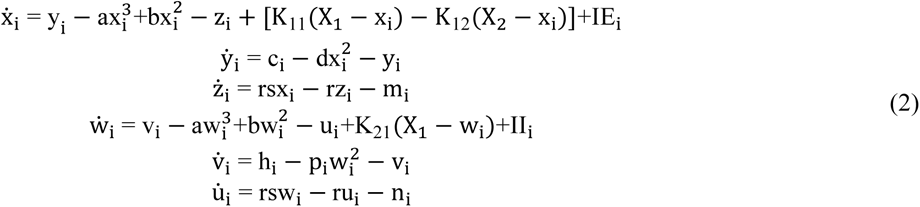

***Eq. 2*.** *Local dynamics of the SJHMR3D model*.

***Legend****: x, y, z, v, w, u – state variables; a, b, c, d – constants representing fast ion channels; r – constant representing slow ion channels; s – bursting strength; IE, II – excitatory and inhibitory input current, respectively; K_11_ – excitatory-excitatory coupling; K_12_ – inhibitory-excitatory coupling; K_21_ – excitatory-inhibitory coupling; X_1_, X_2_ – mean field potential of inhibitory and excitatory subpopulations; m, n, p, h – constant parameters*.

Here, *x(t)* and *w(t)* represent the membrane potentials, *y(t)* and *v(t)* symbolize the recovery variables via fast ion channels, and *z(t)* and *u(t)* the inward currents via slow channels in the membrane for the excitatory and inhibitory populations, respectively. From each simulation, the state variables *x* and *w* have been recorded. Their three modes were averaged to generate the mean field potential of excitatory and inhibitory neurons. The used local parameters for the simulations are described in **Table 1** and initialized with random conditions within certain bounds (see the simulation script for more details).

One of our goals was to create resting-state activity that shows the electrophysiological alpha band frequency oscillation that is present in empirical EEG data. To achieve faster oscillations in the alpha set, we had to not only change the global but also the local parameters. Therefore, we conducted an exploration of the local model and its parameters, which is described in detail in **Supplementary Table 2** and **Supplementary** Figures 1-7. Shortly summarized, we simulated 2250 different combinations of parameters in an uncoupled network (i.e. connection weights = 0) to observe the effect of parameter changes on the local model. From there, we selected those combinations which peaked around 8-12Hz in the spectral density. Due to changes of local parameters, we also changed the noise distribution and integration step size in the second set of simulations (alpha set, **Table 1**).

### Data Analysis

All data preprocessing and analysis was performed with Python and the statistical computing language R. Figures were generated with the Python library Seaborn. To allow the neural signal to stabilize, and to account for random initial conditions in our simulation, we deleted the first 2s of each simulated time series. The BOLD signal was simulated using a mixture of gamma kernels as a hemodynamic response function^44^. To account for the 20s length of the kernel, we removed the first 20s of the BOLD signal. For better comparison of simulated to empirical data, we down-sampled the simulated BOLD signal to a 2Hz sampling frequency. Simulated FC was derived by calculating the Pearson correlation coefficient between all brain regions’ pairs of simulated BOLD signals. Correlation was also used to quantify similarities between the simulated and the empirical FC.

When we mention the ‘optimal fit’ of parameters in the following, we refer to the optimal parameter combination to achieve the highest possible correlation coefficient between the empirical and the simulated FC matrices (empirical-to-simulated FC fit). However, we define the expression ‘optimal working point’ as the parameter set that not only results in the best empirical-to-simulated FC fit but also presents bimodality in the power distribution of the alpha frequency band and reaches the maximum correlation with the FCD matrix (details explained below). We examined whether a relation exists between optimal parameters (global coupling and conduction speed) and graph-theoretical metrics of the SC (**Supplementary Methods**).

### Empirical-to-Simulated FC Fit: Subject Specificity

So far, we compared simulated to empirical FCs of the same subject to optimize our parameters. An important feature of TVB is to provide subject-specific simulations. Since structure shapes function in the brain^29^, one could assume that by providing a subject-specific input, the output fits best to its own empirical data. This assumption would mean that using individual SCs as a structural backbone of a simulation should generate data closer to its own empirical counterpart than to other subjects’ SCs or FCs. To test for subject specificity of our TVB model predictions, we compared all simulated FCs of a subject to the empirical FCs (and SCs) of all other subjects to determine whether the best fit is achieved with the subject’s own FC (SC) or the FC (SC) of another subject. We therefore correlated the set of 105 (1510) simulated FCs in the delta (alpha) set of each subject to all subject’s 50 empirical FCs. **Fig. 2A** shows a schematic overview of these comparisons based on two example subjects. The set of simulated FCs of each subject was compared with the empirical FCs (SCs) of all other subjects, resulting in a highest correlation coefficient for each comparison with the corresponding combination of global coupling and conduction speed parameter values. These results were written in two matrices, the ‘highest correlation matrix’ and the ‘optimal (G,v) parameter matrix’ (**Fig. 2A**). There was no significant linear relation between our optimal parameters and the age of a subject (**Supplementary** Fig. 8-9).

We also quantified whether the underlying SC or the empirical FC determines optimal parameters. Therefore, we use the eta squared effect size^45^, which is computed as the ratio of sum of squares between groups and total sum of squares, grouped here either by the same underlying SC (**Fig. 2C, upper panel**) or by the same empirical FC (**Fig. 2C, lower panel**). This metric has values between zero (no variance explained by the grouping variable) and one (variance completely explained).

### Bimodality: Hartigan’s Dip Test

For every neural signal, we calculated a time-frequency analysis using SciPy’s spectrogram function. The frequency with the highest power values in the spectrogram was then used to test for bimodality. Bimodality can be described as the switching of the neural signal between two modes: one mode with high and another with low amplitude to no oscillation. We used Hartigan’s dip test (as implemented in R) on the distribution of power coefficients of the selected frequency to assess whether power is unimodally or bimodally distributed^46^. With p<0.05, we can reject our null hypothesis that the distribution is unimodal and, therefore, it must at least be bimodal. Bimodality was assessed of the mean simulated neural signal, where the average over the neural signals of all regional time series was taken. We also conducted this analysis on a regional level to account for potential regional differences in bimodality. Examples of a unimodal and a bimodal power distribution are displayed in **Supplementary** Fig. 13.

As the frequency of the oscillations varies across simulations, we did not assess the same frequency for every simulation. Rather, we tested the frequency that is most dominant in the signal, i.e., the frequency that has the highest power in the spectrogram. Therefore, subsequent figures assessing bimodality for a certain parameter range have corresponding figures showing the tested dominant frequencies (e.g. **Fig. 1B and C**).

### Functional Connectivity Dynamics

Since our local SJHMR3D model is highly non-linear, we expected to observe dynamical changes in functional connections as well^47^. It is known that FC is not static over time and rather switches between different modes during, for example, rest^48^, tasks^49^ and learning^50^. To characterize FCD, we required a longer simulation to be able to divide the time series into meaningful time windows. To this end, we conducted 22 minutes simulations for four sample subjects (sex/age: f/30, f/47, m/20, m/73). We used local parameters and global coupling values from the alpha set. Conduction speed was set to the subject-specific optimum derived from the previous analysis of the alpha-set simulations. For each simulated BOLD signal, we created an FCD matrix by correlating numerous FCs from sub-windows of the time series. Time windows of size 60s with an overlap of 40s were used for all FCD matrices. The fit between empirical and simulated FCD was assessed with Kolmogorov-Smirnov distance.

We plan to publish all code after publication. For the reproducibility of our study, we prepared an accompanying data descriptor entitled “Connectomes, simultaneous EEG-fMRI resting-state data and brain simulation results from 50 healthy subjects”^51^, where we present the used multimodal empirical data in raw and processed format as well as the simulation results. This large comprehensive empirical and simulated data set is annotated according to the openMINDS metadata schema and structured following Brain Imaging Data Structure (BIDS) standards for EEG and MRI as well as the BIDS Extension Proposal for computational modeling data. This is the first data set of its kind since it is the first data set that uses the new Brain Imaging Data Structure (BIDS) extension proposal guidelines for computational neuroscience data (https://zenodo.org/doi/10.5281/zenodo.7962031), which is especially designed in an effort to make computational neuroscience studies reproducible. Our code includes a reproduction notebook, which loads the necessary files from the prepared data structure and reproduces the dominant frequency results of an example subject. After publication, the metadata will be available on the EBRAINS Knowledge Graph (https://search.kg.ebrains.eu/).

## Acknowledgements

We gratefully acknowledge the Gauss Centre for Supercomputing e.V. (www.gauss-centre.eu) for funding this project by providing computing time through the John von Neumann Institute for Computing (NIC) on the GCS Supercomputer JUWELS at Jülich Supercomputing Centre (JSC). Additionally, computation has also been performed on the High-Performance Cluster for Research of the Berlin Institute of Health. PR acknowledges Digital Europe TEF-Health 101100700, EU H2020 Virtual Brain Cloud 826421, Human Brain Project SGA2 785907; Human Brain Project SGA3 945539, ERC Consolidator 683049; German Research Foundation SFB 1436 (project ID 425899996); SFB 1315 (project ID 327654276); SFB 936 (project ID 178316478; SFB-TRR 295 (project ID 424778381); SPP Computational Connectomics RI 2073/6-1, RI 2073/10-2, RI 2073/9-1; PHRASE Horizon EIC grant 101058240; Berlin Institute of Health & Foundation Charité, Johanna Quandt Excellence Initiative; ERAPerMed Pattern-Cog, the Virtual Research Environment at the Charité Berlin – a node of EBRAINS Health Data Cloud, Horizon Europe: EBRAINS 2.0 (101147319), Virtual Brain Twin (101137289).

## Author Contributions

P.T. and P.R. conceived the project and designed the experiments. M.S, P.T. and P.R. performed the data acquisition and brain imaging data preprocessing. P.T. and J.M. analyzed the data. J.Z., L.S., D.R., A.S., V.J., G.D., M.B., M.S., A.R.M. and P.R. contributed to the interpretation of the data and the simulation results. P.T., J.M. and P.R. drafted the manuscript. J.M. substantively revised the manuscript. All authors contributed to the preparation of the manuscript.

