## Supplementary information for "Fifty shades of The Virtual Brain: Converging optimal working points yield biologically plausible electrophysiological and imaging features"

### Supplementary Methods

This supplementary section contains a table of region labels (**Supplementary Table 1**), the detailed local parameter exploration for the alpha set (**Supplementary Table 2**), an animation of the bimodality mechanism (**Supplementary Video 1**) and an animation for the fit of functional connectivity, functional connectivity dynamics and bimodality across the global coupling parameter range (**Supplementary Video 2**). We also provide a brief explanation for the difference in regional bimodality between the two sets of simulations, and report statistics on the relation between simulated brain activity and age of the subject.

***Supplementary Table 1: Region labels.*** *Labels for the 68 regions, using the same order as in the connectome. The prefixes "lh_..." and "rh_..." specify left and right hemisphere, respectively.*

| Number | Region label | Number | Region label |
|---|---|---|---|
| **1** | lh_bankssts | 35 | rh_bankssts |
| **2** | lh_caudalanteriorcingulate | 36 | rh_caudalanteriorcingulate |
| **3** | lh_caudalmiddlefrontal | 37 | rh_caudalmiddlefrontal |
| **4** | lh_cuneus | 38 | rh_cuneus |
| **5** | lh_entorhinal | 39 | rh_entorhinal |
| **6** | lh_fusiform | 40 | rh_fusiform |
| **7** | lh_inferiorparietal | 41 | rh_inferiorparietal |
| **8** | lh_inferiortemporal | 42 | rh_inferiortemporal |
| **9** | lh_isthmuscingulate | 43 | rh_isthmuscingulate |
| **10** | lh_lateraloccipital | 44 | rh_lateraloccipital |
| **11** | lh_lateralorbitofrontal | 45 | rh_lateralorbitofrontal |
| **12** | lh_lingual | 46 | rh_lingual |
| **13** | lh_medialorbitofrontal | 47 | rh_medialorbitofrontal |
| **14** | lh_middletemporal | 48 | rh_middletemporal |
| **15** | lh_parahippocampal | 49 | rh_parahippocampal |
| **16** | lh_paracentral | 50 | rh_paracentral |
| **17** | lh_parsopercularis | 51 | rh_parsopercularis |
| **18** | lh_parsorbitalis | 52 | rh_parsorbitalis |
| **19** | lh_parstriangularis | 53 | rh_parstriangularis |
| **20** | lh_pericalcarine | 54 | rh_pericalcarine |
| **21** | lh_postcentral | 55 | rh_postcentral |
| **22** | lh_posteriorcingulate | 56 | rh_posteriorcingulate |
| **23** | lh_precentral | 57 | rh_precentral |
| **24** | lh_precuneus | 58 | rh_precuneus |
| **25** | lh_rostralanteriorcingulate | 59 | rh_rostralanteriorcingulate |
| **26** | lh_rostralmiddlefrontal | 60 | rh_rostralmiddlefrontal |

| | | | |
|---|---|---|---|
| **27** | lh_superiorfrontal | 61 | rh_superiorfrontal |
| **28** | lh_superiorparietal | 62 | rh_superiorparietal |
| **29** | lh_superiortemporal | 63 | rh_superiortemporal |
| **30** | lh_supramarginal | 64 | rh_supramarginal |
| **31** | lh_frontalpole | 65 | rh_frontalpole |
| **32** | lh_temporalpole | 66 | rh_temporalpole |
| **33** | lh_transversetemporal | 67 | rh_transversetemporal |
| **34** | lh_insula | 68 | rh_insula |

***Supplementary Table 2. SJHMR3D parameter exploration.** Parameter values used for exploring the dynamics of the SJHMR3D model.*

| Noise | K11 | n=K12/K11 | μ | σ |
|---|---|---|---|---|
| 0.1 | 0.5 | 0.4 | 2.2 | 2.2 |
| 0.01 | 1.0 | 1.0 | 2.6 | 2.6 |
| 0.001 | 1.5 | 1.6 | 3.0 | 3.0 |
| | 2.0 | | 3.4 | 3.4 |
| | 2.5 | | 3.8 | 3.8 |
| | 3.0 | | | |
| | 3.5 | | | |
| | 4.0 | | | |
| | 4.5 | | | |
| | 5.0 | | | |

***Supplementary Video 1. Mechanisms of bimodality in the alpha range.** Each row shows signals of a single simulation with one global coupling value G. Global coupling G was varied from top to the bottom row with values of [0, 0.025, 0.0328, 0.04]. Conduction speed was set to 100 mm/ms and the other parameters were as in the alpha set. Note: In the video "xi", "eta", "tau" and "alpha" correspond to state variables x, y, z and w of our local model.*

***Supplementary Video 2. Functional connectivity and functional connectivity dynamics in dependence of global coupling.** For subject #2, we show empirical FCD and FC in the left column. The right column animates how simulated FCD and FC change in relation to global coupling. The shift of global coupling is highlighted by the black vertical line in the lowest plot. In the range of ≈0.029, the simulation achieves a good fit according to all three metrics, i.e. bimodality, FCD and FC comparison.*

***Parameter exploration***

To gain insight into the dynamics of the local model, we first conducted a brute-force search with an uncoupled network. We used an SC with 3 nodes and 0 connectivity weights in all connections for our simulations. With no inter-nodal connections, dynamic behavior of the model was driven solely by local parameter variations. All possible combinations of reported local parameters and corresponding values as shown in **Supplementary Table 2** were used for this exploration. The parameters of interest were the dispersion of noise (*D*), excitatory to excitatory coupling ($K_{11}$), excitation-inhibition balance (*n*), the standard deviation ($\sigma$) and mean ($\mu$) of the membrane excitability considered in the neural population. The values and step width of the SJHMR3D parameters were chosen following the original article describing the model (1), where significant differences in mean field amplitude, as well as behaviors such as synchronization of spikes/bursts or chaotic regimes across these parameter ranges were shown. Simulation length for parameter exploration, where we simulated the neural signal, was 5 s. Our aim was to generate alpha oscillations. We performed a spectral density estimate using a periodogram on each generated neural signal. From there, we picked those parameter combinations that showed a peak in the alpha frequency band to investigate further. We refer to **Supplementary Fig. 1** for a summary of local parameter exploration.

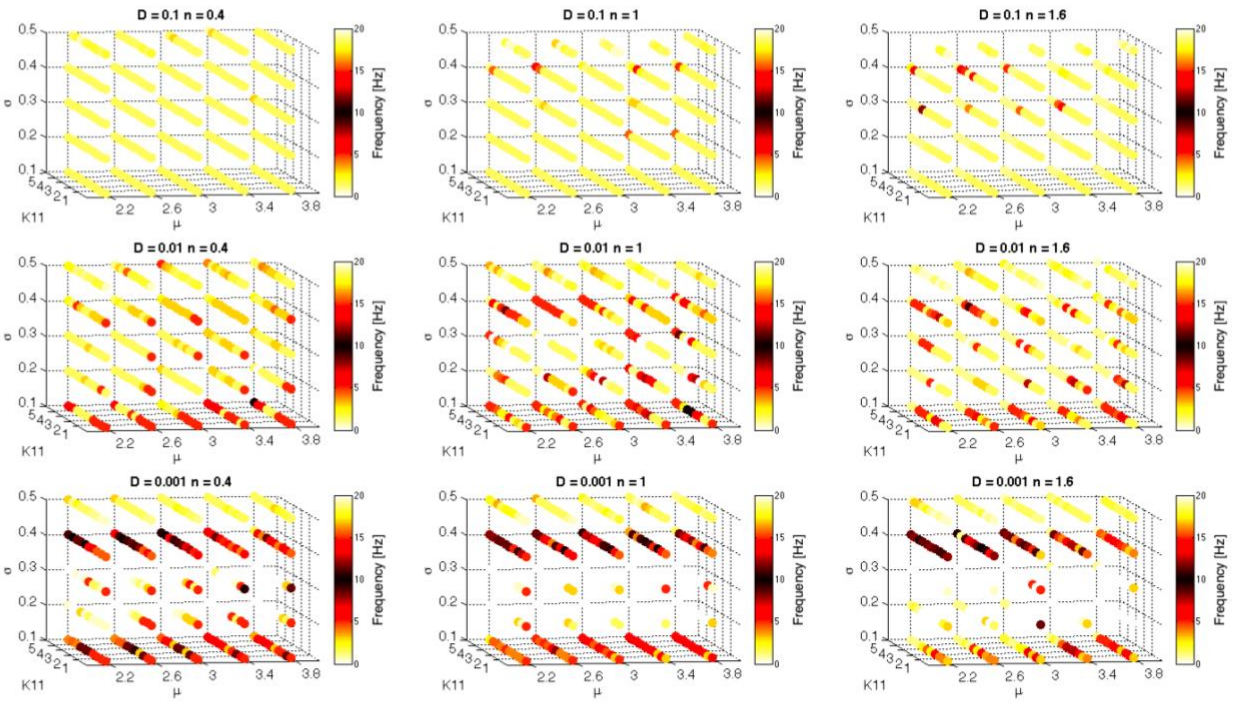


***Supplementary Figure 1. Exploration of excitation/inhibition balance, noise dispersion, mean and standard deviation of population membrane excitability.*** *Each row of scatter plots shows a different noise level. Each column depicts a different value of* $n = K_{12}/K_{11}$. *In each scatter plot, x, y and z axes correspond to* $\mu$, $K_{11}$ *and* $\sigma$, *respectively. Dark colors correspond to alpha band activity. The plots demonstrate that alpha band frequency (dark red) is mostly likely to occur at low noise levels,* $\sigma = 0.4$, *and for many values of* $\mu$, $K_{11}$ *and n.*

In the following, we describe the reasoning for our choice of local parameter values in the alpha set of simulations:

**Noise (*D*):** An appropriate noise value was determined according to the TVB documentation, advising that sensible values of noise dispersion are at approximately < 1% of the dynamic range of a model's state variables. Alpha band frequency was observed at noise dispersion = 0.001. In some cases where noise dispersion = 0.1, the simulations were unstable and generated NaN values, particularly when $\sigma$ and $n$ were high.

**$\sigma$:** Alpha band frequency was observed for values of the standard deviation of the membrane excitability in range of 0.4. The signal makes a transition through a complex oscillatory behavior, continuous spiking, bursting in an alpha band frequency, and slower bursts. In **Supplementary Fig. 2**, the plots show time series for different parameters.

**$K_{11}$:** The bursting behavior was ubiquitous for $\sigma = 0.4$. Frequency increased slightly with $K_{11}$. This is consistent with $K_{11}$'s role as excitatory-excitatory coupling. Peak frequency was observed at 11 Hz for $K_{11} = 4$ (**Supplementary Fig. 3**).

**$\mu$:** With increasing the mean of the membrane excitability, the power in the alpha band decreases (**Supplementary Fig. 4**). Therefore, we selected $\mu = 2.2$, where alpha band power was highest. The relationship of frequency, $\mu$ and $K_{11}$ is displayed in **Supplementary Fig. 5**.

The local dynamics were then coupled with the rest of the network, whereby we used the full SC of the first subject and examined parameter values in the context of the global dynamics. In the global model, we examined $n$, the ratio of inhibitory and excitatory coupling (= $K_{12}/K_{11}$). With all other parameters set to values defined in **Supplementary Table 1**, and global coupling = 0.03, we tested all three $n$ values in a coupled model using the SC of the first subject. In **Supplementary Fig. 6**, one can see that for 3 tested values of $n$, there was a peak at the alpha frequency in the periodogram, with the greatest alpha power observed at $n = 0.4$.

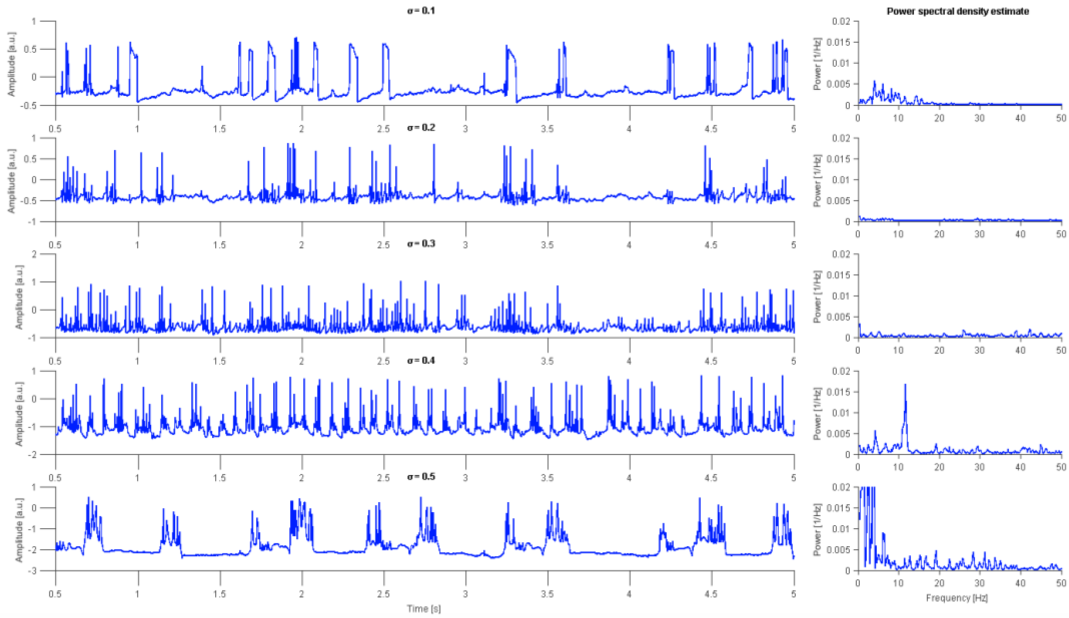


***Supplementary Figure 2. Exploration of standard deviation of membrane excitability (σ).*** *Each row of plots shows the timeseries of the mean field potential (MFP); On the right, the power spectral density (PSD). For* $\sigma = 0.4$*, the power in the alpha band is the highest. Parameters from the exploration that are not mentioned here are the ones used in the alpha set.*

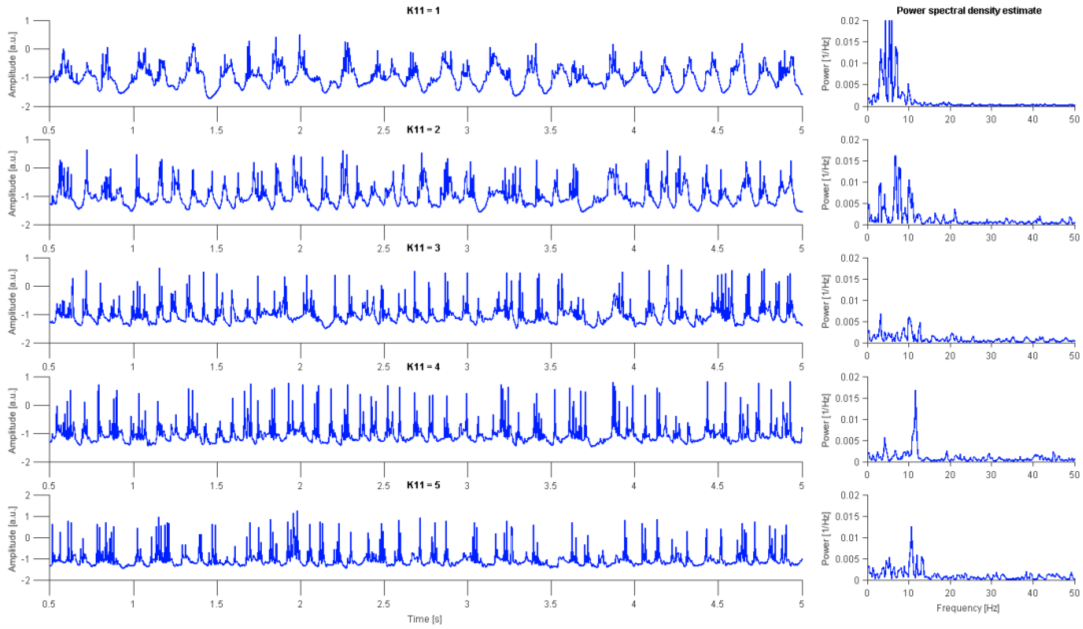


***Supplementary Figure 3. Exploration of excitatory to excitatory coupling (*$K_{11}$*).*** *Each row of plots shows the time series of the MFP. On the right, the PSD. For* $K_{11} = 4$*, the power in the alpha band was highest.*

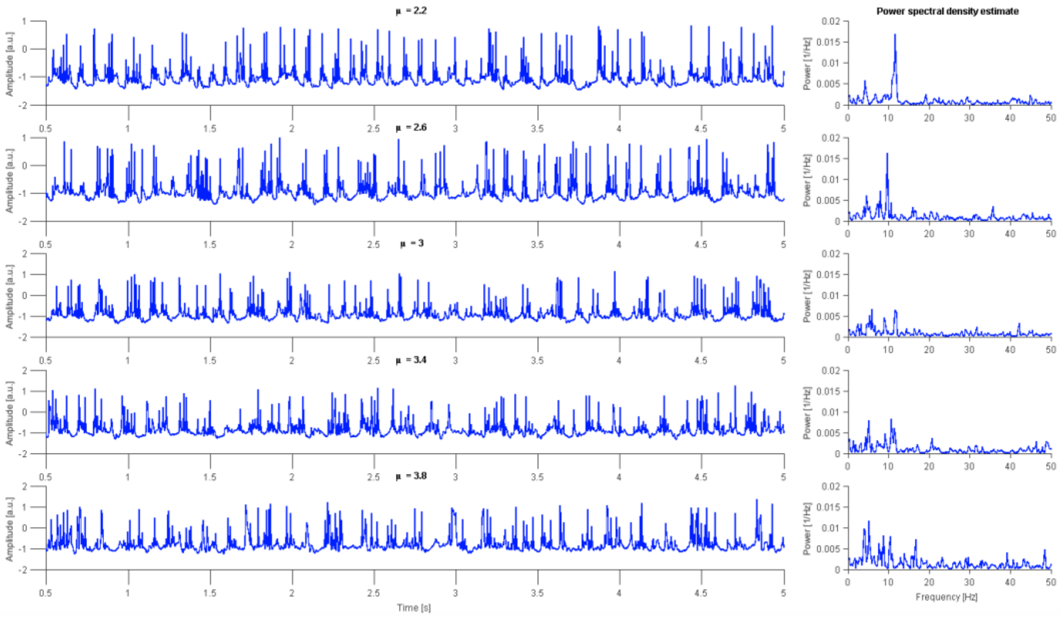


***Supplementary Figure 4. Exploration of the mean of membrane excitability (μ).*** *Each row of plots shows the time series of the MFP. On the right, the PSD. For μ = 2.2, the power in the alpha band was highest.*

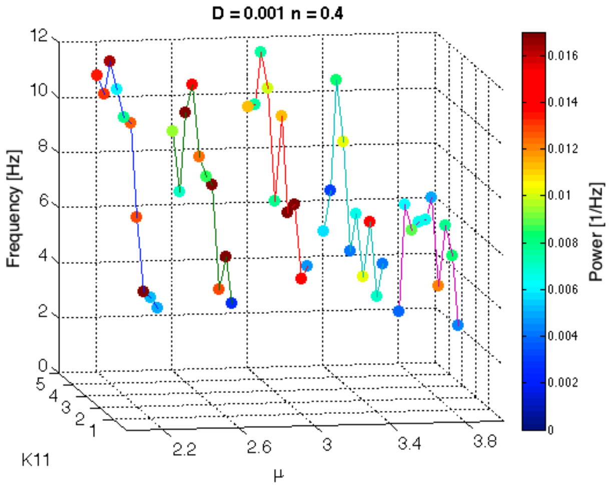


***Supplementary Figure 5. Alpha band frequency can be observed for nearly all μ, and high $K_{11}$ values.*** *However, the power (indicated by color) is highest for a low μ. Lines are drawn for better identification of points.*

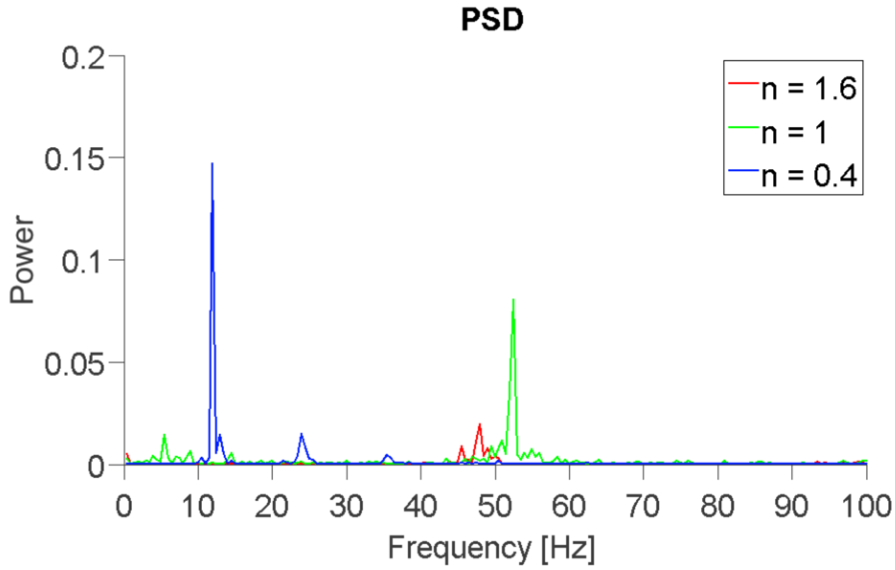


***Supplementary Figure 6. Power spectral density (PSD) estimate for different values of n.*** *The desired alpha band frequency is observed only for n = 0.4. All coupled simulations are performed with global coupling G = 0.03.*

***Difference in regional metrics***

We observed that bimodality was differentially distributed across brain regions in our delta set simulations. The presence of bimodality closely correlated with nodal degree. The stronger the node was connected, the more likely it was that this node's simulated activity showed a bimodal power distribution. This effect was not apparent in the alpha simulations. With the aim to understand why alpha set simulations do not show regional bimodality differences, we showed that regional variation in oscillatory behavior is highly dependent on local parameters. Thus, minute differences in local parameter values may be responsible for whether we observe regional variation in bimodality in the alpha and delta set. In our example, we show regionally varying dependency of frequency on $K_{12}$ (inhibitory to excitatory coupling). That is, the value of $K_{12}$ influences how regions with varying nodal degree exhibit particular frequency values. In **Supplementary Fig. 7**, we show that for low values of $K_{12}$ (i.e. $K_{12} = 0.1$), frequency is higher among regions with low degree; in contrast, frequency is low for regions with high degree. Higher degree means a greater influence of other regions on the nodal oscillations; this influence is mostly dampening to the nodal oscillations as few of the other regions oscillate. Increasing $K_{12}$ even more (i.e. $K_{12} = 1$) and more of the low degree regions begin to oscillate faster. At $K_{12} = 1.6$, the whole global system switches into a highly oscillating state. Increasing $K_{12}$ further to $K_{12} \approx 4$, the low degree regions' oscillations start to slow down again. At $K_{12} \approx 6.6$, all regions again show a high frequency oscillation. Increasing $K_{12}$ beyond a certain point ($K_{12} > 9$) results in instability of the system (NaNs). In short, we aim to show how local parameters drive regional variation in frequency, in an effort to understand why the delta set, but not the alpha set, shows regional differences in bimodality.

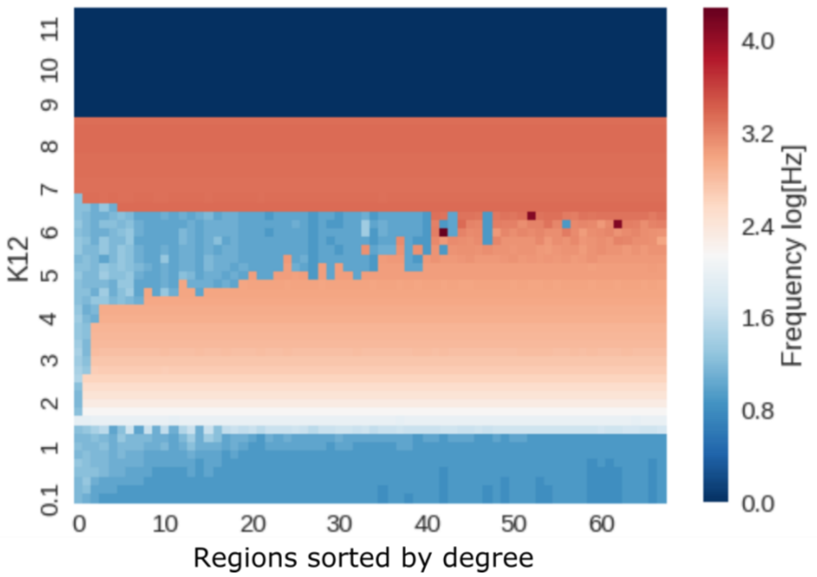


***Supplementary Figure 7. Impact of inhibitory to excitatory coupling strength ($K_{12}$) on the coupled network.*** *Regions were sorted by their degree in ascending order. Frequency was log-scaled to better visualize regional differences in the slow frequency band. All other parameters were set to the alpha set parameters defined in Table 1, with global coupling set to G=0.032 and conduction speed to v=100mm/ms. For increasing coupling, the global network observes different states of synchronized and unsynchronized oscillations. In the range of $K_{12} \in [4, 6]$, oscillation frequency increases with degree.*

***Influence of subject age on simulation outcomes***

There was no significant linear relation between our optimal parameters and the age of a subject (**Supplementary Fig. 8**). Within the alpha set, the relation of age to optimal coupling almost reached the significance level of 0.05. Subsequently, Partial least squares (PLS) analysis was used to reveal patterns in SC and simulated FC related with age (2). SC and FC matrix were averaged across rows, that is FC/SC was averaged across all connections of a node, and resulting subject-specific vectors were concatenated across all subjects to form two matrices – one for SC and one for simulated FC. A third SC-FC matrix was generated by computing the correlation between a region's SC and FC vectors. The three matrices were stacked and used as input to a behavior PLS (3). PLS decomposes the correlation between brain features and age into sets of orthogonal latent variables (LVs). Each of them explains a different aspect of the relation. Significance and reliability of the estimation was assessed using permutation and bootstrapping resampling. We found one significant LV ($p < 0.0001$) where SC, FC and SC-FC were all reliably predictive of age (**Supplementary Fig. 9**). The correlation was strongest for the simulated FC ($r = -0.568$) but still reliable for the other two features (SC, $r = -0.372$ and SC-FC, $r = -0.373$). Region-wise loadings are shown in the horizontal bar plot. Regions above the threshold (= 2.0) have a reliable age-related effect in SC, FC and SC-FC coupling.

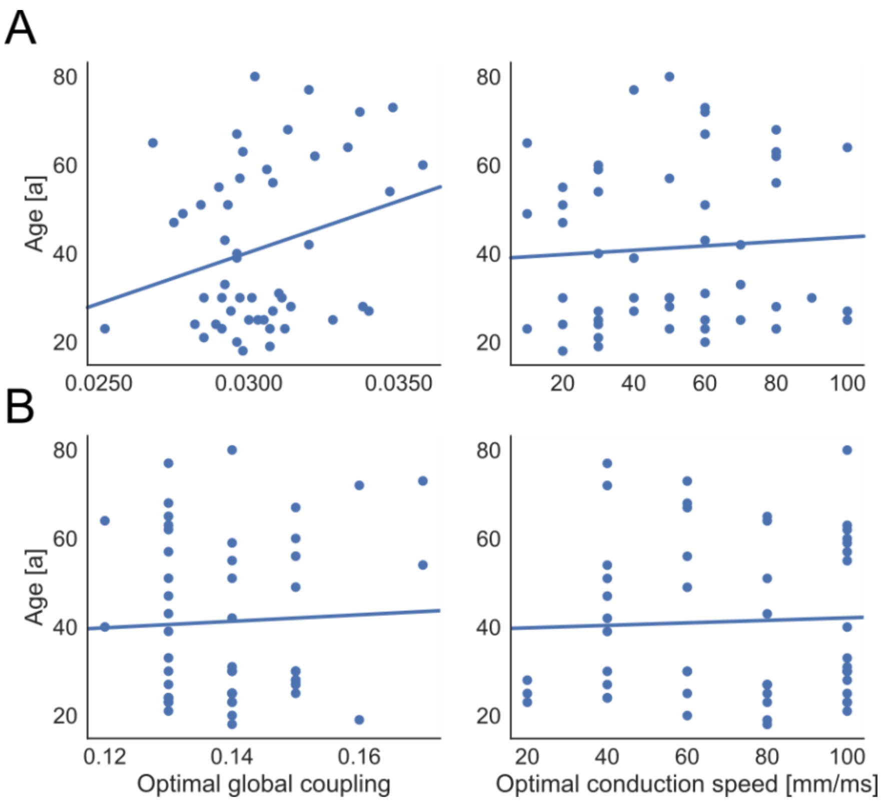


***Supplementary Figure 8. No significant relation between the optimal parameters and age.*** *For the alpha (A) and delta (B) set, the plots show age against the optimal parameter of each subject. The relation of age and optimal global coupling in the alpha set was* close to significant with r = 0.2659, p-value = 0.062. The other p-values were > 0.6 for all comparisons.

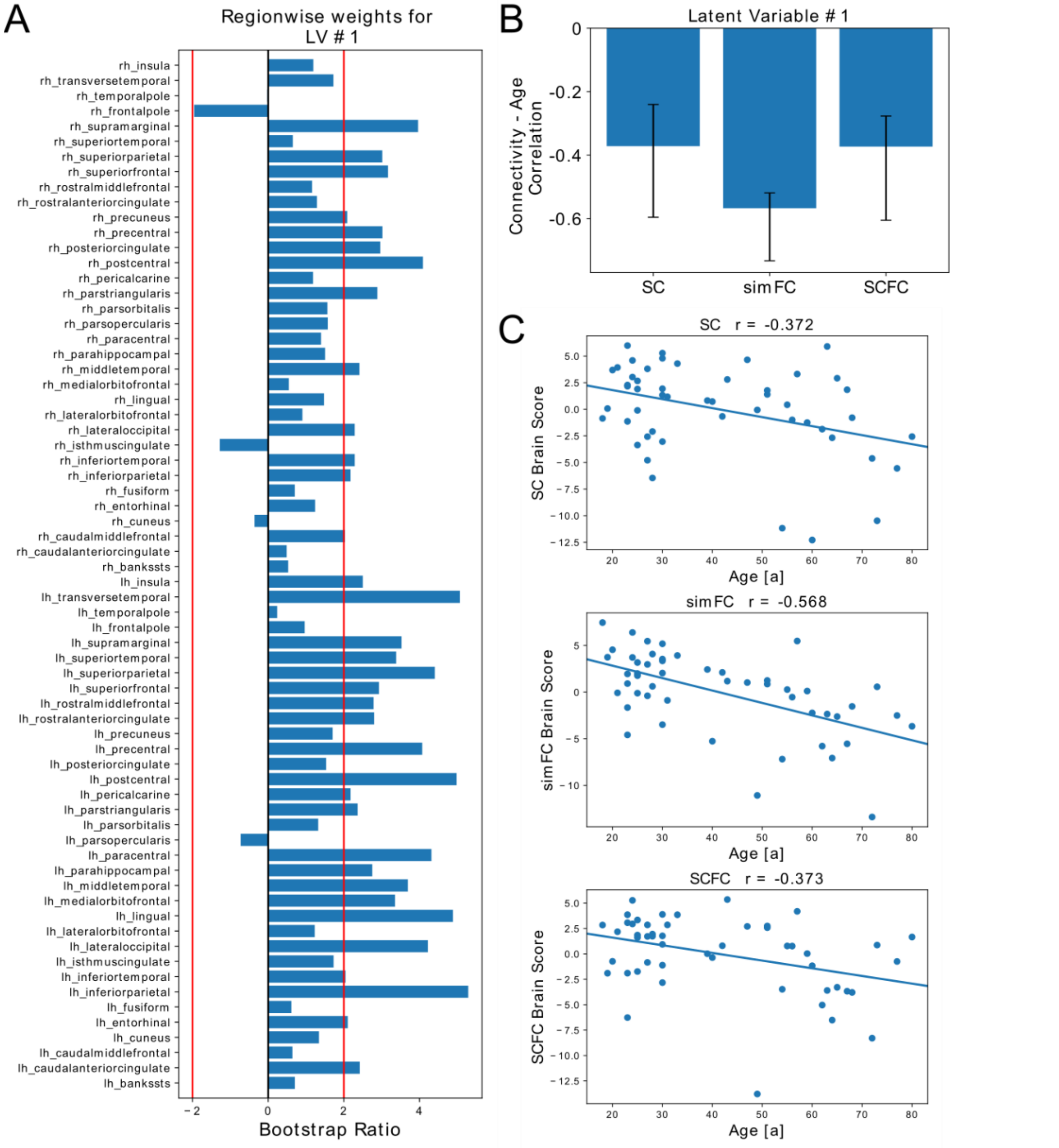


***Supplementary Figure 9. PLS analysis to reveal a possible relation between age and SC as well as simulated FC.*** *From the PLS analysis only the first latent variable (LV) was found to be significant and is reported here. This analysis uncovers an inverse relation between the SC, simulated FC (simFC), and SC-FC metric with age. All three measures decrease with increasing age. A: PLS bootstrap ratios are shown for the LV loadings. Regions above the threshold (= 2) have reliable age-related reductions in SC, FC, and SC–FC coupling. B: Bar graph showing the correlation between SC and age, simFC and age, and SC–FC coupling and age. Confidence intervals on each of these correlations are shown. There is a reliable relationship between each of these and age. C: Scatterplots visualizing the correlations between SC brain scores and age, simFC brain scores and age, and SC–FC brain scores and age.*

***Mechanisms of bimodality in the alpha range (Supplementary Video 1)***

Bimodality in our model occurs as a consequence of the global inhibition that results from the negative baseline of state variable *x* representing the membrane potential of the excitatory population. With increasing global coupling *G* (top to bottom row), inhibition is strengthened, leading to global synchronization, bimodality and finally, oscillation death. In the main part of this article, we viewed the state variables of the SJHMR3D model as scalars, when in fact they are vectors with a length of three. The elements represent the activity of different modes of the neural mass model. The model is a mean field reduction of a neural network of 1000 coupled Hindmarsh Rose neurons. Mode decomposition was used to capture the behavior of different neuron clusters into three modes. For the detailed analysis in the following video, we focus on the activity of the different modes. We show how the global input to a node impacts the local dynamics and creates a bimodal power distribution.

***Description of the video:*** Each row shows signals of a single simulation with one global coupling value *G*. Global coupling *G* was varied from top to the bottom row with values of [0, 0.025, 0.0328, 0.04]. Conduction speed was set to 100 and the other parameters were as in the alpha set. Note: In the video "xi", "eta", "tau" and "alpha" correspond to state variables *x, y, z* and *w* of our local model, respectively.

Columns from left to right:

1. Time-series of state variable "*x*" of the left lateral occipital area in our network.
2. The phase space of the excitatory populations first mode from the left occipital area. It shows how the three state variables of the excitatory subpopulation evolve over time.
3. The excitatory phase plane of the first mode of one node. State variables "x" and "y" build the axis of the plane. Red arrows indicate the direction of the flow at a specific position. Four colored (red, green, blue and black) example trajectories were drawn into the phase plane to highlight the existence of limit cycles for certain values of *z, w* and global input. The three stars "*" show the values of "x" and "y" of the three excitatory modes for a given timepoint. Because of their movement, we can see whether the system evolves along the limit cycle or settles in a fixed point. The phase plane was drawn for static values of "z", "w" and the input derived from the global network at the timepoint shown in the plot of the first column.
4. Input to the left lateral occipital area through our global network.
5. Distribution of power coefficients for the dominant frequency.

Rows from top to bottom:

1. The activity of a node with a global coupling *G = 0*, i.e. no connection between regions. Global network input to the selected node is shown in the fourth column. In the first three columns, we observe the local behavior, which shows a square-wave bursting, a state where the neuron fires a group of spikes followed by a quiescent period. In the phase space and phase plane we see behaviour similar to a fold and homoclinic bifurcation (which has been already described at a single Hindmarsh-Rose neuron (4)). Whenever the neurons are bursting, we see a limit cycle in the phase plane. When the system returns to rest, all trajectories in the phase plane tend towards a fixed point. For this simulation we can see a unimodal distribution of power at 4.9 Hz, which corresponds to the frequency of the bursts.
2. Synchronized activity at a global coupling of G = 0.025. We see spikes, i.e. single excitations, in the time series of state variable x. These spikes can be seen as "bursts" with a single (or sometimes up to three) spikes. If we look to the global input plot, we can see when all regions synchronize their spikes, the global input is positive (up to a value of 2). But most of the time the global input is negative, resulting in global inhibition. However, the local excitation (given by parameters "μ" and "σ" of the SJHMR3D model) is strong enough to overcome this inhibition and generate spikes. The power of variable x in log-log coordinates is unimodally distributed at a frequency of 15.2 Hz.
3. Bimodal activity at a global coupling of 0.0328. The global negative input (inhibitory) is strong enough to suppress any activity for a certain period. However, when global input rises (due to activity in other regions with lower degree) it allows spikes in the node. This can also be seen in the phase plane. Most often, the trajectories tend towards a fixed point. However, when global inhibition is reduced, we can observe a limit cycle and a spike in the time-series. This gives rise to periods with oscillation and periods without oscillations – resulting in a bimodal power distribution at 9.2 Hz.
4. Oscillation death at a global coupling of 0.04. No neural activity is observed. The trajectories tend towards a fixed point at all times.

***Supplementary Table 1: Parameters of optimal working point distributions.***

| | **delta set** | | **alpha set** | |
|---|---|---|---|---|
| | **global coupling** | **conduction speed [mm/ms]** | **global coupling** | **conduction speed [mm/ms]** |
| **median** | 0.14 | 80 | 0.03 | 50 |
| **mean** | 0.1398 | 70.8 | 0.0304 | 50 |
| **standard deviation** | 0.0113 | 26.56 | 0.0021 | 25.0713 |

***Empirical-to-Simulated FC Fit: Relation with Graph-Theoretical Measures***

We examined whether a relation exists between optimal parameters (global coupling and conduction speed) and features of the SC. We checked for the relation to numerous graph-theoretical measures like average degree, characteristic path length or global clustering coefficient of SC (weights and tract lengths) and FC. In both simulation sets, optimal coupling decreased with increasing average degree, $r = -0.68$ ($p < 0.0001$) for delta and $r = -0.88$ ($p < 0.1 \cdot 10^{-15}$) for alpha set. This result is because the output of a node is multiplied by weights of the SC and then scaled by the global coupling factor (**Eq. 1**). Therefore, strongly connected SCs require lower couplings than weakly connected ones. We found no significant correlation between global coupling and any other graph-theoretical measures or conduction speed and any graph-theoretical measures, neither for the FC nor the SC.

We found that optimal coupling decreased with increasing average SC weights. We explain this result as follows: To realistically model local dynamics within one node, a certain amount of input from other nodes is needed. Therefore, if the SC weights are low, a sufficient high global coupling value is necessary to scale up the input from other nodes. A link between coupling and global efficiency in the SC has previously been reported[1], but the association to mean SC has not yet been explicitly made.

***Empirical-to-Simulated FC Fit: Subject Specificity***

Individual fits (where simulated FC and empirical FC are from the same subject) are not higher than all-to-all fits (where simulated FC and empirical FC come from different subjects). Some empirical FCs correlate highly with all simulated FCs, as can be observed by the (red) horizontal stripes in the empirical FC-simulated FC plot (**Fig. 2B and Supplementary Fig. 10A, left**). That is, the goodness of fit is more determined by specific features in the empirical FC connectome than the simulated FC (and the underlying SC). Future analysis should further investigate which features are of importance here.

Analyses of empirical FC versus empirical FC correlations and SC versus SC correlations (**Supplementary Fig. 10B, left and right,** respectively) showed that empirical FCs were less homogeneous than the SCs across subjects as displayed by the overall higher correlations among the SCs (colorbar limits of both plots). Individual SC and empirical FC (**Supplementary Fig. 10B, middle**) did not exhibit subject-specific fits.

There is a linear relation between the fit of empirical FC to SC and the fit of simulated to empirical FC (independent of which subject, **Supplementary Fig. 10C, left**, $r = 0.73$ ($r = 0.8$) with $p < 0.001$ ($p < 0.001$) for delta (alpha) set).

Correlations between empirical SC and FC, empirical SC with simulated FC and simulated and empirical FC are shown in **Supplementary Fig. 10C (right)**. The distribution of highest correlations between empirical FCs and our simulations is shown there as the red density estimate. We compared the alpha and delta sets and found that correlations of the fits were in the same range of values (Wilcoxon signed-rank test on the distribution of highest correlations of the two sets was non-significant). For both sets, we observed that the correlation between individual SC and empirical FC (green) is on average lower than that of simulated to empirical FCs (red). In other words, by using only SC, we cannot predict the empirical FC as good as by using the SC in a simulation, which points to the added value by TVB. Although simulated FCs fit best to their specific underlying SC among all other SCs, they still fit better to their empirical counterpart (blue < red, **Supplementary Fig. 10C, right**). Thus, the combination of SC and our TVB model is more like an FC than an SC.

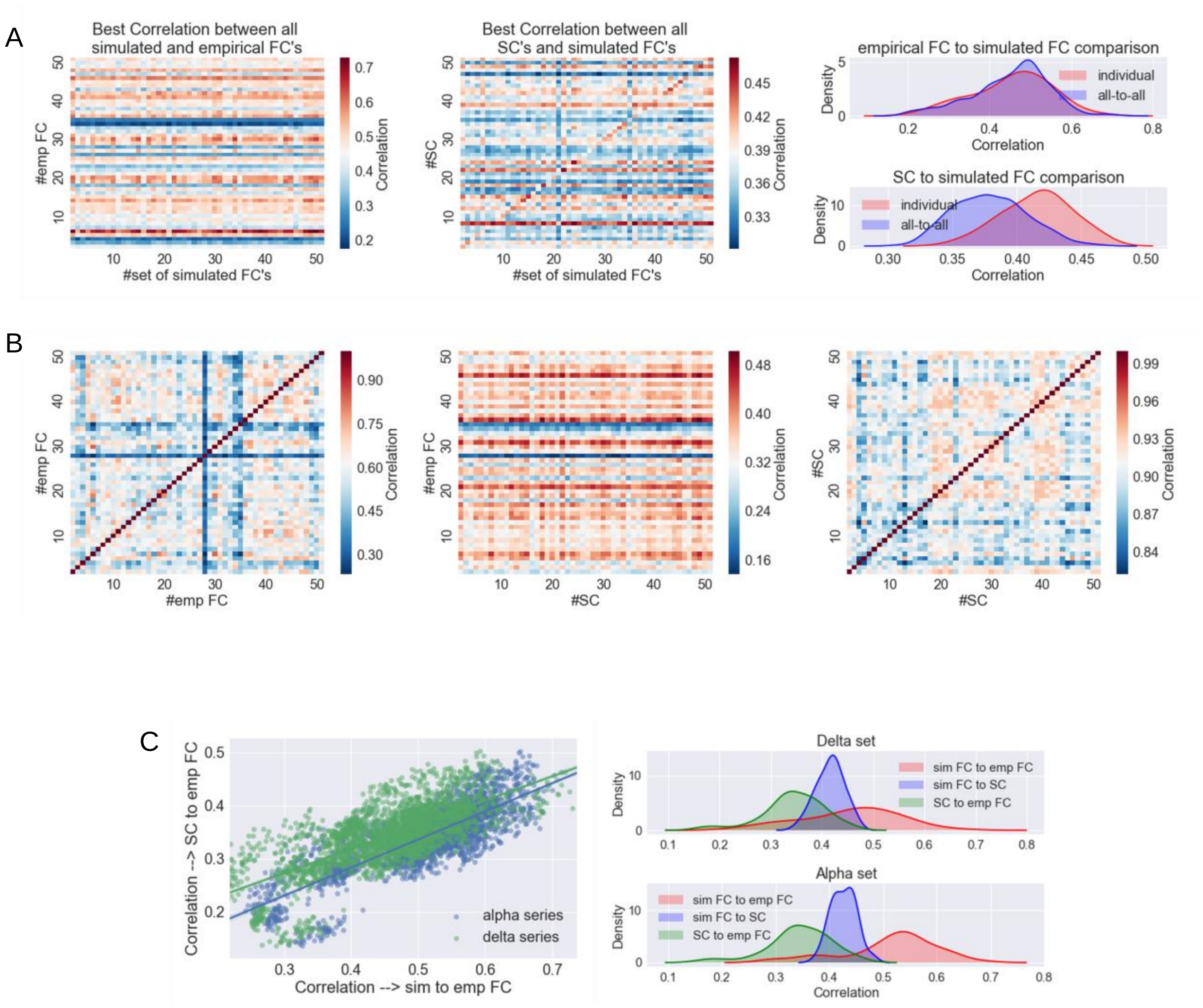


***Supplementary Figure 10. A:*** *Delta set: Left: We correlated all 105 simulations of one subject to all 50 empirical FCs. For each empirical FC that we correlated to, there was one best (i.e. highest) correlation. In other words, comparing all simulations from one subject to all FCs, generated 50 highest possible correlations. This process resulted in a 50x50 matrix when comparing for all 50 subjects ('highest correlation matrix', Fig. 1A). The entry (i,j),* $1 \le i, j \le 50$*, in this*

*matrix represents the highest correlation based on the comparison of the empirical FC matrix of subject i with the set of all simulated FCs of subject j. Along the diagonal of the matrix, we found the highest possible correlation for the individual subject prediction (i.e. predicting a subject's empirical FC by using the same subject's SC for simulation). Horizontal red and blue stripes already indicate that some empirical FCs can be predicted better than others, regardless of the SC used for the simulation. Middle: The same as the left matrix but now comparing set of simulated FCs to SCs, instead of empirical FCs. We tested the possible prediction of the SCs from the simulation. A vague red diagonal indicates a best individual fit, pointing to the fact that the simulated FC contains typical features of the used SC. Right: To test for subject specificity, we took the diagonal values (i.e. subject's individual prediction) and tested their distribution against the values above and below the diagonal (i.e. all-to-all prediction). Comparing simulations to empirical FCs showed no significantly better individual prediction. Individual SCs can be predicted significantly better than all-to-all prediction (two-sample Kolmogorov-Smirnov test* $d = 0.4963$ *(*$d = 0.5492$*) and* $p < 0.1 \cdot 10^{-9}$ *(*$p < 0.1 \cdot 10^{-12}$*) for delta (alpha)). This result indicates how strongly the SC shapes the simulated FC. All of your density plots were smoothed with a Gaussian kernel.* ***B****: Left matrix shows correlation between all 50 empirical FCs, e.g. the FC of subject #27 is very different to the rest FC (blue horizontal and vertical line). Right matrix is the same for SCs, i.e. a direct comparison between the SCs of our data set. The SCs seem to be much more homogeneous (compare the high correlation coefficients on the colorbar). Middle matrix shows how similar FCs are to SCs. Already without simulations, some FCs can be predicted better by many SCs (e.g. red horizontal line for subjects #29 and #35).* ***C****: Left: Scatterplot of the correlation coefficients based on the comparison of any SC to empirical FC against the highest correlation that can be achieved between empirical FC and the set of simulated FCs. The better an empirical FC correlates to any given SC, the better we can predict it by simulating on that SC. Correlation* $r = 0.73$ *(*$r = 0.8$*) with* $p < 0.001$ *(*$p < 0.001$*) for delta (alpha) set. Right: Best possible individual prediction of empirical FC by SC and simulated FC, as well as SC by simulated FC (top: delta set, bottom: alpha set). Simulations correlate on average higher to empirical FC (red) than to SC (blue). Also, the prediction of the empirical FC is on average better with simulation rather than using the SC directly (red > green). Though we were not able to reproduce subject-specific FC features, our simulations still compared in general better to functional rather than structural data. All of our density plots were smoothed with a Gaussian kernel.*

***Supplementary Table 2: Eta squared.*** *Explained variance of optimal parameters by empirical FC and the simulations underlying SC for both sets.*

| | **delta set** | | **alpha set** | |
|---|---|---|---|---|
| **explained variance** | **by underlying SC** | **by empirical FC** | **by underlying SC** | **by empirical FC** |
| **for global coupling** | 0.4629 | 0.2826 | 0.7968 | 0.0175 |
| **for conduction speed** | 0.0735 | 0.0913 | 0.0989 | 0.0809 |

Regions with numbers 4 and 38 are the entorhinal cortices of both hemispheres showing nearly no bimodality (**Supplementary Fig. 11C and D, left**). With a degree of 7.5 and 6.4 for left and right hemisphere, respectively, the entorhinal cortices share the lowest degrees of the whole connectome. Their activity seems to be mostly uncoupled from the rest of the system.

***Alpha Bimodality: Regional Differences***

We also analyzed bimodality and the frequency of the neural signal of each region individually. Results are visualized in **Supplementary Fig. 11** with data averaged across subjects. The plots were additionally averaged over either the whole range of conduction speed values (**Supplementary Fig. 11A and C**) or global coupling values (**Supplementary Fig. 11B and D**). Averaging across subjects and global coupling dimensions (**Supplementary Fig. 11B and D)** renders the p-values of the Hartigan's dip test of bimodality non-significant. Therefore, we displayed the percentage of subjects with bimodality (i.e. $p > 0.05$) on the colorbar for all plots (**Supplementary Fig. 11, left**). Optimal values of conduction speed occur at the higher end of the range. The trend of more subjects with bimodality around G = 0.14 (0.0295) and at conduction speeds of 60-100 mm/ms (> 20 mm/ms) in the delta (alpha) set, is indicative of the optimal parameter space that shows significant bimodality within individual subjects. For the same parameter combinations, oscillations in the respective frequency band were generated (**Supplementary Fig. 11, right**).

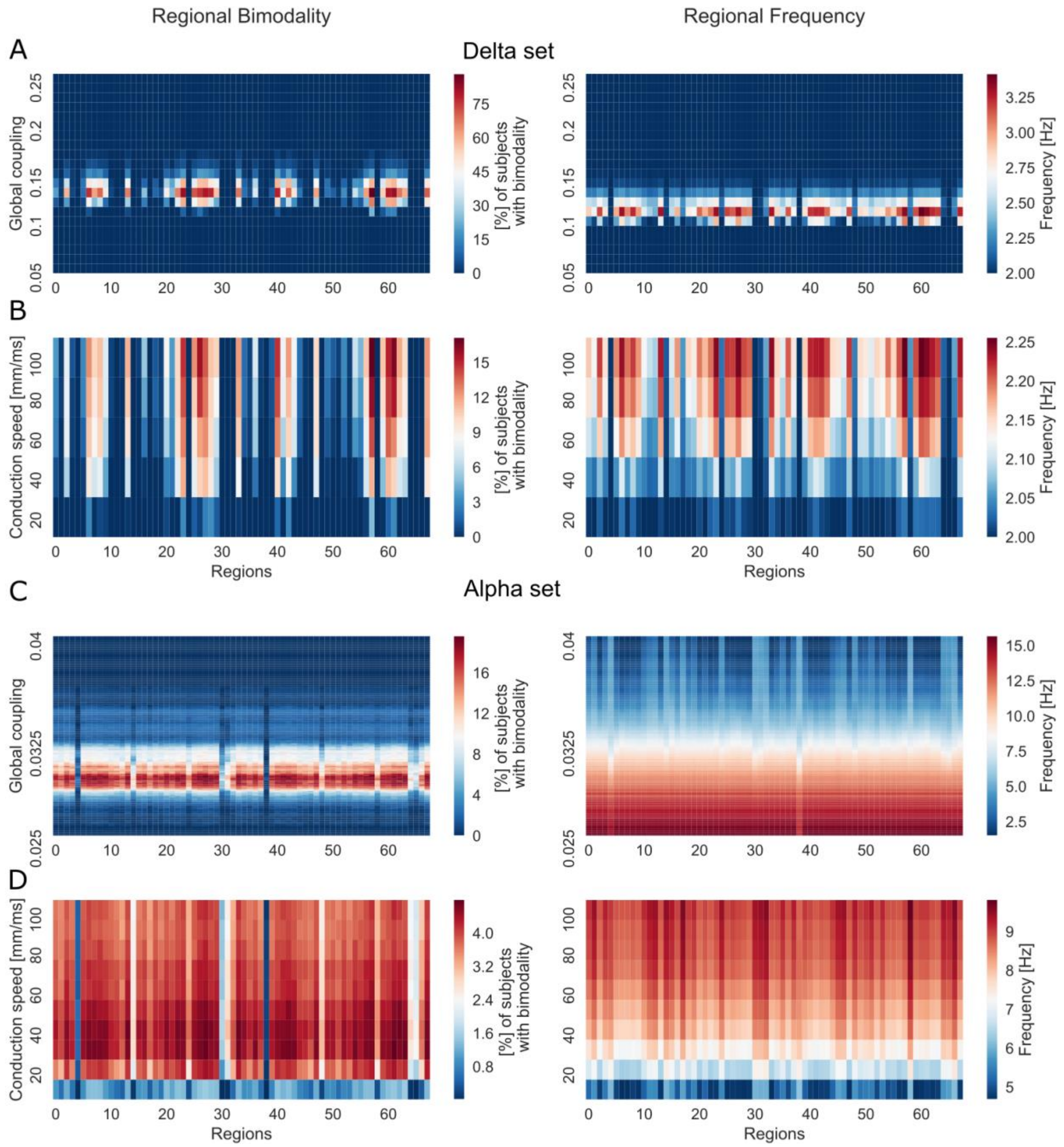


***Supplementary Figure 11. Regional behavior of neural signal for delta and alpha set simulations.*** *Panels A and C depict the effect of global coupling on regional bimodality and frequency (averaged over all subjects and the whole range of conduction speed values). Bimodality and fast oscillations only occur at a critical range of global coupling. The peak in bimodality occurs at coupling values slightly higher than for the maximum frequency. Panels B and D display the effect of conduction speed on regional bimodality and frequency (averaged over all subjects and the whole range of global coupling values). The oscillations become faster with higher speed, but bimodality is observed only for speeds greater than 20 mm/ms.*

Within the delta set, we found an interesting distribution of features across regions, with only some showing bimodal power distributions and faster oscillations (**Supplementary Fig. 11A and B**). In order to understand the basis of these inter-regional differences, we examined the relationship between degree and bimodality. We took the regional Hartigan's dip test p-value at optimal parameter points (based on the highest correlation of the empirical-to-simulated FC fit) and plotted it against the average degree of each region. We show that regions with a bimodal power distribution were strongly connected within the SC network (**Supplementary Fig. 12C).** Regional bimodality in anatomical space is shown in **Supplementary Fig. 12B**. Bimodality is observed primarily along the parietal and temporal cortex ($p < 0.05$ for most of these regions). However, not all subjects at their optimal parameter combinations show bimodality within these regions (these are the outliers in **Supplementary Fig. 12A**). At their optimal parameter point, 80% (92%) of subjects in the delta (alpha) set possess bimodal power distribution for their mean neural signal.

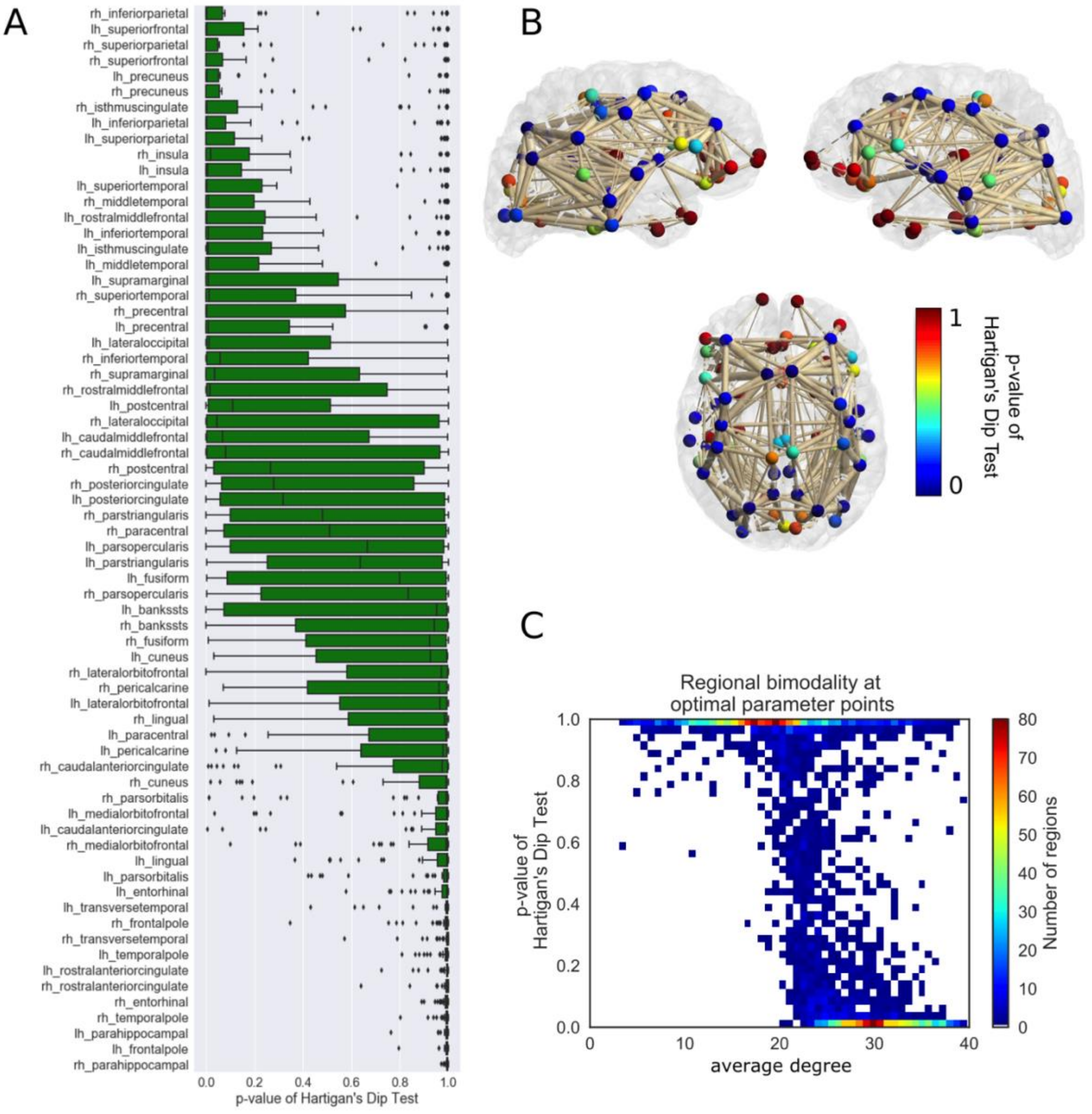

***Supplementary Figure 12. Anatomical distribution of bimodality in the delta set. A:*** *At the optimal parameter point of each subject, we assessed bimodality for all regions on the left (lh_...) and right (rh_...) hemisphere. Distribution of Hartigan's dip test p-values per region are shown in a boxplot. Regions are sorted from top to bottom by decreasing average degree.* ***B:*** *3D graph of our brain network model. To display edges, we used the mean SC across subjects thresholded to a sparsity of 17% (i.e. showing only 17% strongest connections). Edge thickness represents the SC weight. Node colors represent the mean p-value of the Hartigan's dip test over all subjects at optimal parameter points, with blue colors indicating bimodality.* ***C:*** *2D histogram showing relation between regional bimodality and mean degree. At the optimal parameter combination, we assessed bimodality in each region. All graphs indicate how strongly connected brain regions present bimodality at the optimal parameter point.*

***Mechanisms Underlying Alpha Bimodality***

With the present local parameters (**Table 1**) and no global coupling, we observed a bursting behavior in the neural signal that has been documented by the authors of the model[3]. A fold/homoclinic bifurcation is characteristic for this type of burst, which is sometimes called square wave bursting, where the trajectory follows a limit cycle in the 3D phase space and generates spikes. Meanwhile, the slow variable ($z$) increases and moves an unstable fixed point towards the limit cycle, generating a saddle homoclinic bifurcation, i.e. the unstable fixed point touches the cycle and breaks it up. The trajectory then switches towards a stable fixed point, which corresponds to the resting period. During "rest", the slow variable decreases and moves the unstable system back towards the stable fixed point creating a fold bifurcation - a transition from rest back to spiking.

When state variable $x$ - reflecting the membrane potential of the excitatory population - was coupled with a linear scaling function to other regions of the network, it results in an inhibiting effect during baseline activity (when its values were negative) and an excitatory effect during spikes (when its values were positive).

For even higher couplings ($G = 0.025$), we observed a synchronization of local dynamics with global input. The regional neural signal displayed single bursts with few spikes rather than the bursting behavior we had observed at $G = 0$. In the phase space, there were only a few turns on the limit cycle before the homoclinic bifurcation occurs. The global input became even more negative (i.e. -4) than at $G = 0.01$. We showed that the spiking in the regional neural signal actually coincided with the times at which the global input reached zero. This result can be interpreted as a temporary reduction in global inhibition that leads to spikes in the local model.

For very high global coupling (*G = 0.04*), oscillation death occurred and very slow noisy fluctuations in the neural signal were observed. The neural signal and the global input had negative amplitudes and fluctuated very little. The trajectory remained at a stable fixed point. The state of bimodality occurred in-between the synchronized and the oscillation death state (*G = 0.032*). Namely, we observed a switching between oscillation death and spikes that occurs in synchrony with a reduction in global inhibition. We also showed this behavior in **Supplementary Video 1.**

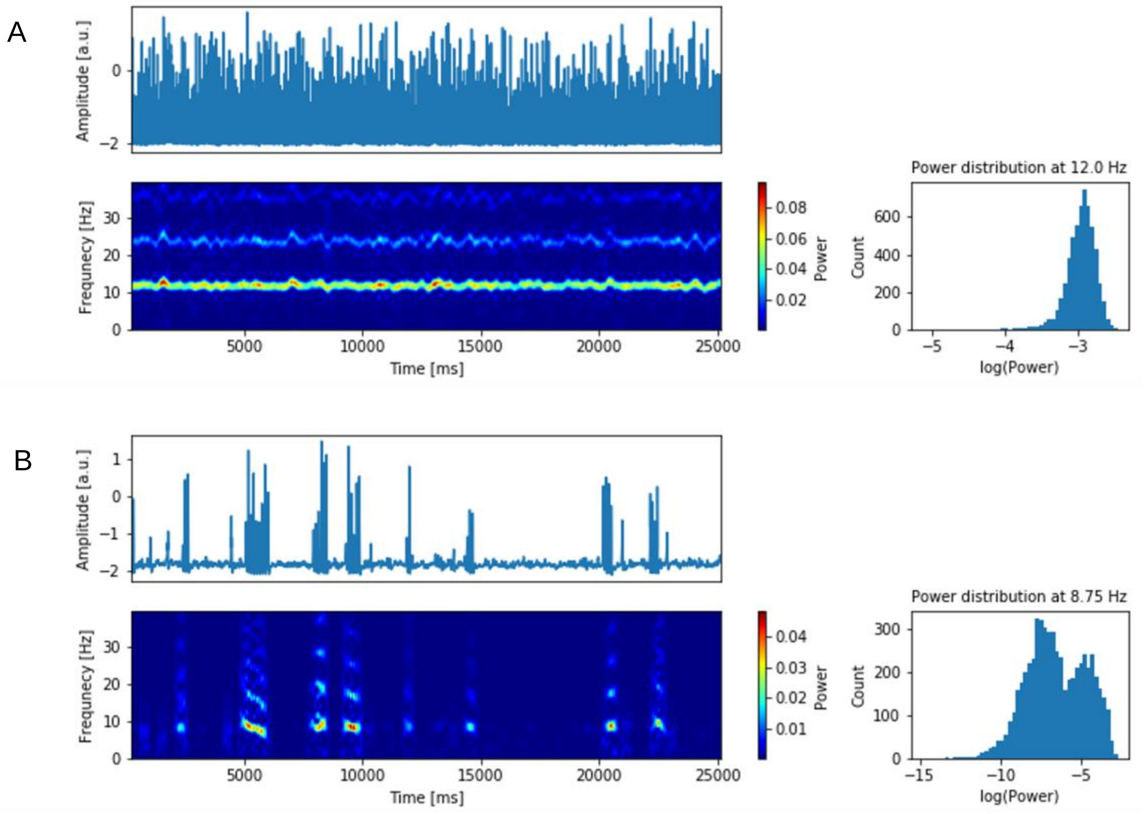


***Supplementary Figure 13. Schematic of subject specificity analysis and example of a unimodal and a bimodal power distribution of the simulated neural signal.*** *Time series were simulated using parameters as in the alpha set with different global coupling strengths G = 0.03 and G = 0.0328 for panels A and B, respectively, and conduction speed equal to 100mm/ms in both cases. Each panel depicts the neural signal (upper plot), a time-frequency analysis (lower plot) and, on the right, the distribution of power coefficients at the highest power frequency for a unimodal (bimodal) signal in panel A (B). In panel B, the time series and spectrogram show a switching between high- and low-amplitude oscillations. Hartigan's dip test p-values are 1 and < 0.001 for the unimodal (panel A) and bimodal (panel B) distribution, respectively.*

***Age and sex distribution in the used data set***

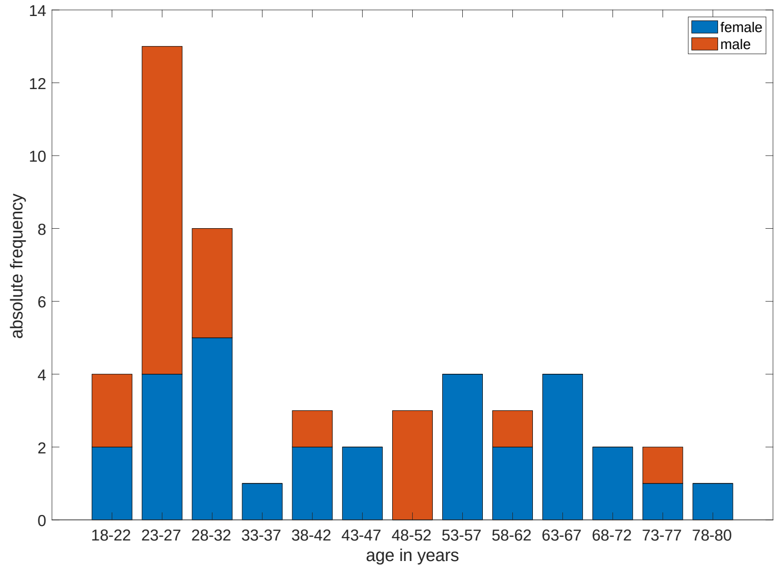


***Supplementary Figure 14: Age and sex distribution in the data set.***

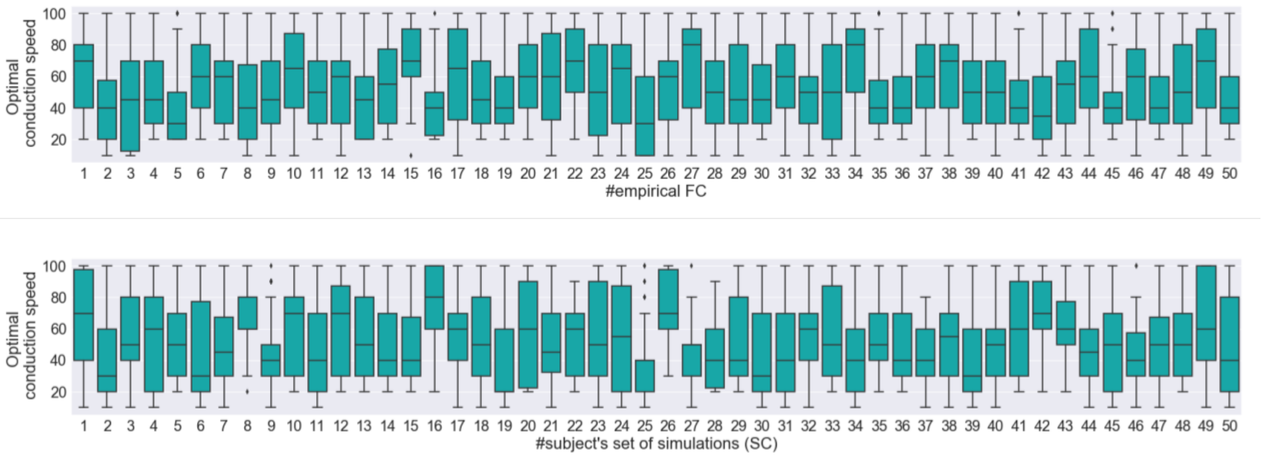


***Supplementary Figure 15:*** *Optimal parameters for conduction speed (results displayed for the alpha set)**:** We analyze how the optimal parameter distributions change between subjects. The parameters are influenced by the underlying SC as visible in the upper panel. Each boxplot displays the optimal conduction speed values for each subject (best reproducing empirical FCs of all subjects). The variance of optimal conduction speed for a given subject's SC is similar to the variance of the same across individual subjects' SCs. In the lower panel, we display the influence of the empirical FC on the distributions of optimal conduction speed values. In contrast to optimal global coupling being determined by the SC, conduction speed is neither determined by the SC nor by empirical FC**.***

***Supplementary Table 5: Sex differences in optimal working point distributions.*** *We found no significant differences between male and female participants' optimal working points distributions. P-values are based on a Student's t test* *between the mean values of male vs. female participants.*

| | delta set | | | alpha set | | |
|---|---|---|---|---|---|---|
| | **Male** | **Female** | **p-value** | **Male** | **Female** | **p-value** |
| **Mean optimal conduction speed** | 71.0 | 70.7 | 0.97 | 46.5 | 52.3 | 0.43 |
| **Mean optimal global coupling** | 0.1405 | 0.1393 | 0.73 | 0.0302 | 0.0306 | 0.51 |
